# Nanometer-scale RNA protein clusters (RPCs) Foster Helicase Activity of DEAD-box eIF4A

**DOI:** 10.64898/2026.05.23.727435

**Authors:** Him Shweta, Masaaki Sokabe, Nancy Villa, Christopher S. Fraser, Yale E. Goldman

## Abstract

DEAD-box RNA helicases are central regulators of RNA metabolism, employing ATP-dependent mechanisms to remodel RNA structure and RNA-protein interactions, yet how helicase catalysis is coordinated with multi-subunit interactions between RNA and protein remains unresolved. Translation initiation helicase, eukaryotic initiation factor 4A (eIF4A), which acts as an intrinsically non-processive enzyme, is essential for unwinding structured mRNAs, relies on cofactors to achieve physiological activity. Here we uncover an unexpected RNA-helicase state of eIF4A, demonstrating that eIF4A forms nanometer-scale RNA-protein clusters (RPCs) of ∼2-5 MDa in presence of its physiological cofactors eIF4B and eIF4G, RNA and ATP under near-physiological concentrations. Using a single molecule approach, we directly resolve the formation of discrete clusters that recruit multiple copies of proteins with RNA upon ATP addition and show that RPC formation correlates with helicase activity *in vitro*. Further, we find eIF4B as a key determinant of this multi-subunit assembly. Its intrinsically disordered regions (IDRs) together with structured RNA-recognition motifs (RRMs) drive multivalent RNA-dependent clustering, critical for efficient helicase activity. Disrupting eIF4B-RNA interactions through a targeted point mutation (F139A) in the RRM reduces both the cluster size and the helicase activity, further establishing a functional link between cluster formation and catalytic activity. Consistent with these findings, in-cell diffusion measurements reveal markedly slower diffusion of wild-type eIF4B compared with the RNA-binding-deficient mutant, indicative of RPC formation within the cellular environment. Together, our results reveal regulated helicase clustering as a previously unrecognized characteristic of the translation initiation machinery, linking ATP-dependent DEAD-box helicase activity to nanometer-scale RNA-protein clusters and translation initiation regulation.

## Introduction

ATP-dependent RNA helicases of the DEAD-box family are central and widespread regulators of RNA metabolism^1^. In eukaryotic translation initiation, the helicase eukaryotic initiation factor 4A (eIF4A) is essential for removing secondary structure in the 5′ untranslated region of mRNAs to enable ribosome recruitment and scanning^2^. Although indispensable, eIF4A helicase activity is almost negligible and non-processive on its own, with physiological functional activity accelerating through cooperative interactions with other translation initiation accessory factors^2,3^. Single-molecule, biochemical, kinetic and structural studies have established that eIF4G acts as a scaffolding activator that allosterically stimulates eIF4A, while eIF4B enhances productive RNA engagement and couples ATP hydrolysis to strand separation, together enhancing unwinding activity^4–11^. In line with this framework, a single-molecule optical trap assay showed that eIF4A functions as an ATP dependent processive helicase in presence of eIF4B and eIF4G^10^, defining eIF4A as a cofactor-coupled mechanochemical system rather than an autonomous helicase.

Beyond enzymatic activity, growing evidence indicates that RNA metabolism is influenced by the organizational state of ribonucleoprotein assemblies. Multivalent RNA-protein interactions can generate higher-order assemblies spanning stoichiometric complexes, dynamic nanometer-scale clusters, and larger biomolecular condensates^12–15^. Assembly of these complexes is driven by combinations of complex reversible interactions at nanomolar concentrations, and in some cases above critical micromolar concentration values, liquid-liquid phase separation (LLPS) possibly coupled with percolation^16,17^. Biomolecular nano-clusters and larger condensates extend across a wide range of biomolecules containing intrinsically disordered regions (IDRs)^18^, RNA recognition motifs^19^, prion-like domains^18^, and stress granule associated proteins including FET like proteins, FUS^20^, EWSR1^21^, TAF15^21^, DDX3^22^, DDX4^23^, LAF-1^24^, DDX6^25^ etc. In this framework, eIF4A functions as an ATP-dependent RNA chaperone that suppresses LLPS RNA condensation and limits stress granule formation^26^, whereas eIF4B, through extensive IDRs can self-associate in the micromolar range into large, dynamic oligomers and mesoscale assemblies^27^. A recent discovery shows that the closely related DEAD-box helicases DDX3X and DDX3Y form nanometer-scale RPCs well below the critical LLPS concentration range driven by IDRs at the N- and C-termini of the helicase sequences, and that clustering enhances catalytic activity^22^. This work established clustering as a native and functional organizational state distinct from both monomeric enzymes and LLPS micrometer-sized condensates. Unlike DDX3X and DDX3Y, which contain extended intrinsically disordered regions that can directly promote clustering, eIF4A contains relatively limited disordered regions, raising the possibility that clustering may instead emerge through cofactor-dependent multivalent interactions.

Given the evolutionary conservation of the DEAD-box core and shared reliance on ATP-dependent RNA remodeling and multivalent interactions between RNA- and IDR-driven assembly, these findings raise the possibility that regulated clustering may represent a broader organizational principle among DEAD-box helicases. However, whether cofactor-stimulated eIF4A can adopt RNA-protein cluster states, despite its very limited IDRs and whether the existence and functional relevance of such a state might influence the activity of the eIF4A helicase, has been unresolved.

Here, we test these possibilities using single-molecule fluorescence spectroscopy and in-cell diffusion measurements. We show that human eIF4A alone does not assemble into RNA-protein clusters in vitro, but the addition of RNA, ATP, and its physiological cofactors leads to markedly decreased diffusion rate, indicative of RPC formation. eIF4B mediated multivalency and RNA engagement nucleate cluster formation, whereas eIF4G provides a scaffolding function that may recruit and organizes multiple copies of eIF4A within these RPC assemblies. eIF4A’s ATPase activity dynamically regulates cluster formation and its stability. Together, these findings extend the clustering paradigm established for DDX3X/DDX3Y and other proteins to the translation initiation machinery and identify regulated helicase clustering as a previously unrecognized organizational principle underlying eIF4A function.

The multi-parameter single-molecule confocal fluorescence spectroscopy approach used enables simultaneous extraction of various signals using fluorescently labeled double-stranded RNA (dsRNA) bound to eIF4A, eIF4B, and eIF4G. Here, we tested whether these assemblies represent simple multivalent associations and whether they are functionally coupled to helicase activity. We assessed how RNA engagement regulates cluster formation and helicase activity, defined their composition and stoichiometry, identified the domains of eIF4B critical for nucleating clusters, and evaluated the cellular relevance of RPC formation.

## Results

### Nanometer-scale RNA and cofactor-stimulated eIF4A protein clusters (RPCs)

To capture the physical state of RNA engaged by the translation initiation helicase machinery, and test whether eIF4A forms nanometer-sized RPCs, similar to DDX3X/Y^22^ and FUS^20^, we used fluorescence correlation spectroscopy (FCS) which enables quantification of the diffusion constant of fluorescent particles diffusing through a femtoliter detection volume to infer the apparent hydrodynamic size (see Methods). In combination with eIF4A and its physiological cofactors, eIF4B, and eIF4G, we employed a fluorescently labeled double-stranded RNA (dsRNA) substrate that has been validated for investigating eIF4A helicase activity in earlier studies^28^. The RNA substrate consisted of a 24 base-pair (bp) duplex region with a 20-nucleotide (nt) 5′ single-stranded overhang (Fig. 1A). The longer strand was labeled at its 3′ end with Alexa647, while the complementary strand carried a 5′ Cy3 fluorophore, enabling sensitive detection of the RNA-eIF4A assembly’s state by FCS, fluorescence resonance energy transfer (FRET) and concentration of doubly-labeled particles relative to that of Cy3-only and AF647-only species. Individual dsRNA molecules freely traverse the ∼1 fL confocal observation volume producing discrete fluorescence bursts under pulse-interleaved excitation (PIE) with alternating 532 and 640 nm laser illuminations. The mean occupancy time of each complex within the detection volume (𝜏_D_) is inversely proportional to its diffusion coefficient, *D*, according to 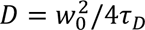, where 𝑤_0_ defines the beam waist radius. By auto-correlating photon streams arising from direct excitation of the Alexa647 over 2.5 min acquisition intervals, we quantified diffusion coefficients of RNA-protein complexes, allowing us to directly link cofactor-stimulated helicase engagement with changes in molecular size and assembly state. The concentration (*C*) of fluorescent particles is given by C = *n/(V·N*_A_) where *n* is the average fluorophore occupancy in the confocal volume (typically 0.3 – 0.4), *V* is the detection volume and *N*_A_ is Avogadro’s number.

**Figure 1.**
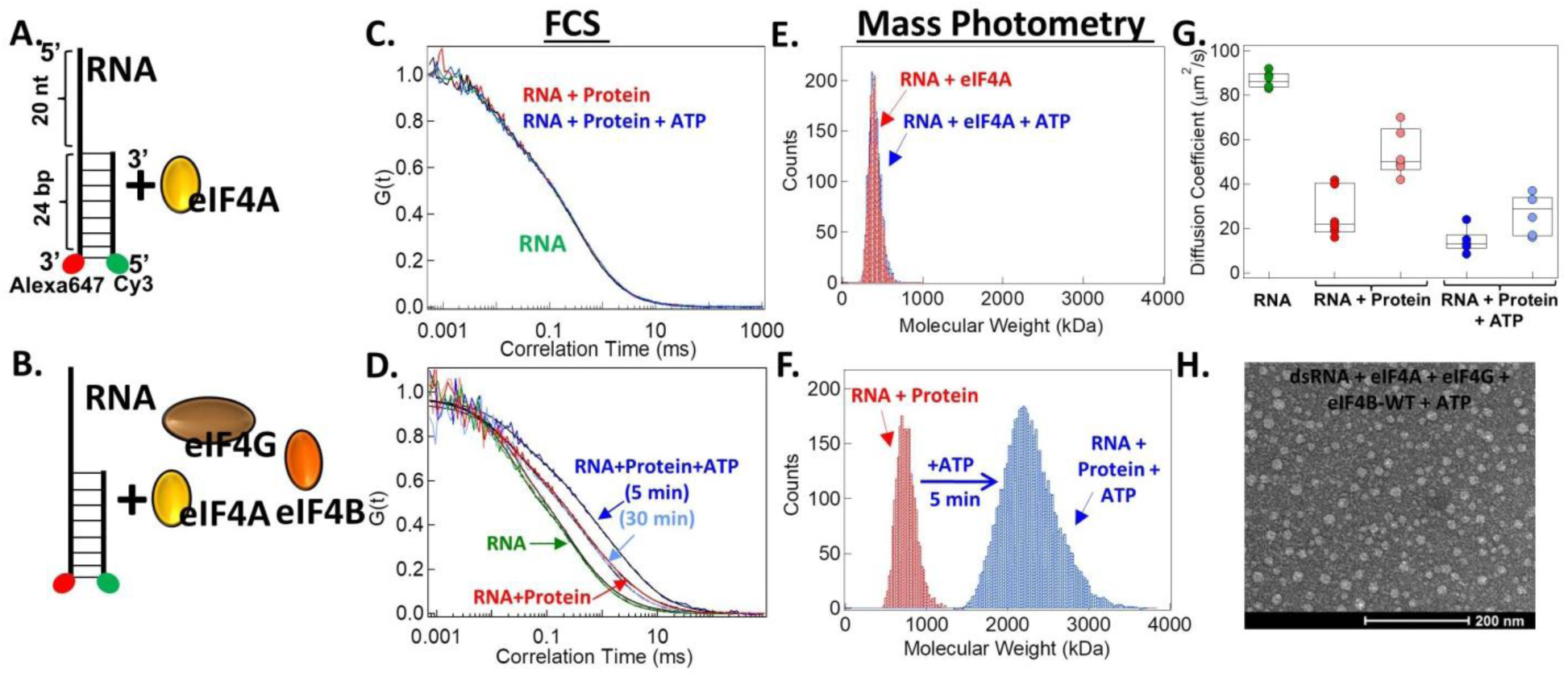
eIF4A form RNA-protein clusters only in presence of the accessory proteins. (A, B) Schematics of Alexa-647 and Cy3 labeled double stranded RNA (dsRNA), and either eIF4A or eIF4A, eIF4B and eIF4G. (C) Normalized auto-correlation curves obtained from the FCS measurements are plotted for 0.5 nM dsRNA only (green), following the addition of 0.5 µM eIF4A (red), and subsequently with 1 mM ATP added plus 5 min of incubation (blue). (D) Normalized auto-correlation obtained from the FCS measurements are plotted for 0.5 nM dsRNA only (green), following the addition of 0.5 µM of eIF4A, eIF4G, eIF4B at 5 min (red) and 20 min (light red), and subsequently 1 mM of ATP at 5 min (blue) and 30 min (light blue). All the fits are shown in black. The dsRNA-eIF4A-eIF4B-eIF4G complex shows slower diffusion relative to dsRNA after ATP addition due to RNA-protein cluster formation. (E) Histogram of the interferometric contrast for dsRNA-eIF4A complex before (red) and after ATP addition (blue). (F) Histogram of interferometric contrast for dsRNA-eIF4A-eIF4B-eIF4G complex before (red) and after ATP addition (blue). (G) Diffusion coefficients (μm^2^/s) of RNA (green), dsRNA-eIF4A-eIF4B-eIF4G complex at 5 min (red), 20 min (light red), dsRNA-eIF4A-eIF4B-eIF4G complex after ATP addition at 5 min (blue) and 30 min (light blue). The box plot displays the mean, 25^th^ percentile (box bottom), and 75^th^ percentile (box top) compiled from 6 independent measurements. (H) Representative electron microscopy image of negatively stained eIF4A, eIF4B, eIF4G (0.5 μM each) with 0.5 nM dsRNA plus Mg-ATP. Scale bar, 200 nm.

dsRNA (0.5 nM) on its own diffused rapidly with a diffusion time (𝜏_D_) = 0.33 ±0.01 ms (mean ± s.e.m., n = 6), consistent with a freely diffusing 27 kDa dsRNA molecule (expected *D* = 86 ±1.45 µm² s⁻¹; (mean ± s.e.m., n = 6)). Addition of eIF4A (0.5 µM), in the absence or presence of 1 mM Mg-ATP, had no measurable effect on dsRNA diffusion (Fig. 1A, C). In contrast, a mixture of dsRNA (0.5 nM) with eIF4A, and cofactors eIF4G and eIF4B (0.5 µM each, Fig. 1B) resulted in a reproducible slowing of diffusion, consistent with binding of a protein complex to the dsRNA (Fig. 1D) increasing the apparent *M*_w_ of the particles (*D* = 26.8 ±4.56 µm² s⁻¹; (mean ± s.e.m., n = 6)). Strikingly, subsequent addition of 1 mM Mg-ATP to this helicase cofactors complex markedly slowed diffusion (Fig. 1D), yielding a diffusion rate of *D* = 14.9 ±1.86 µm² s⁻¹; (mean ± s.e.m., n = 6) and apparent ∼5.2 MDa *M*_w_ of the diffusing complexes. Such a pronounced change in diffusion cannot be reconciled with specific low-stoichiometry RNA-protein binding. Instead, given the inverse relationship between diffusion coefficient and hydrodynamic radius, *r*, according to 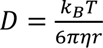, where *k_B_* is Boltzmann constant, *T* is temperature, and *η* is viscosity, and the cubic scaling of mass with radius (𝑚𝑎𝑠𝑠 ∝ 𝑟^3^), this decrease in diffusion rate corresponds to RNA-protein clusters whose apparent molecular masses are increased by approximately 190-fold relative to free dsRNA. Quantitatively, the observed *D* values imply presence of RPCs in the ∼4-5.5 MDa range, far exceeding the mass of individual eIF4A-eIF4B-eIF4G monomers (∼180 kDa), and therefore implying collections of multiple RNA and protein molecules. The apparent molecular weight of these compact assemblies corresponds to a diameter of ∼22 nm.

As a complementary quantitative approach to directly assess the mass distribution of RNA-eIF4A complexes before and after ATP addition, we used mass photometry an optical method to measure molecular weight of the molecules landing on a glass coverslip utilizing contrast arising from optical interference between light scattered by individual particles and the reflection from the glass. The interferometric the contrast scales linearly with molecular weight^29^. Calibration was performed using standards of known molecular weight (Fig. S1A and S1B). The molecular weight estimated by mass photometry for the RNA-eIF4A complex (359 ±35.77 (mean ± s.e.m., n = 4) kDA, Fig. 1E and S1C). Similarly, the dsRNA-eIF4A-eIF4B-eIF4G assembly shows an apparent molecular mass of 872.9 ±78.42 (mean ± s.e.m., n = 6) kDa (Fig. 1F and S1C) before addition of ATP which increased to ∼2.08 ±0.07 (mean ± s.e.m., n = 6) MDa (Fig. 1F and S1C) upon addition of ATP and 5 min incubation, both in excellent agreement with the FCS measured values.

As a third approach, we employed transmission electron microscopy (TEM) on negatively stained samples containing 0.5 nM dsRNA, 1 mM Mg-ATP and either eIF4A alone (0.5 µM) or the fully constituted eIF4A-eIF4B-eIF4G complex (0.5 µM each). Under these conditions, the dsRNA-eIF4A complex failed to yield any discernible higher order structure (Fig. S1D). In contrast, the fully complemented eIF4A-eIF4B-eIF4G complex formed distinct, round, and homogeneous nanometer-scale RPC particles with dsRNA (Fig. 1H). These observations reveal a highly cooperative tendency of cofactor-stimulated eIF4A helicase to drive RNA into discrete, nanoscale clusters, motivating a deeper exploration of the physical and functional nature of this unexpected organizational state.

### RPCs are functionally coupled and crucial to helicase activity

The RNA strand separation activity of eIF4A helicase was quantified by using single-burst fluorescence spectroscopy to detect decrease of FRET efficiency between the Cy3 and AF647 probes and decrease of proportion particles containing both RNA strands. The donor-only (DO), doubly-labeled (DL), and acceptor-only (AO) populations were tracked at 2.5 minute intervals before and after ATP addition. Reduction of the proportion of high FRET (*E* > 0.7) DL particles indicates separation of the strands within an RPC, whereas reduction of the DL population indicates short and long strands escaping RPCs or shifting into DO and AO RPCs. The corrected change in DL, the product of the DL particles and the proportion of high FRET efficiency within the DL component, indicated as DL_HF_ (see Methods) following ATP addition provides readout of the total helicase activity. The reduction of DL is also accompanied by a corresponding increase in the DO and AO populations. After an initial measurement of the dsRNA, protein was added (first dashed line in Fig. 2C and D). No change in the DL population was detected for the RNA-eIF4A complex before and after the ATP addition for another hour (Fig. 2A, C). Upon addition of eIF4A together with eIF4G, eIF4B, a very small decrease of the DL and DL_HF_ populations took place, and then, in sharp contrast to eIF4A alone, a much more pronounced decrease of DL and DL_HF_ occurred within a few minutes upon addition of 1 mM Mg-ATP (Fig. 2B, D), consistent with the onset of helicase activity. The time-dependent decrease of the DL_HF_ population following ATP addition was fitted with an exponential function to extract both the amplitude of DL_HF_ change and the rate of helicase activity.

**Figure 2.**
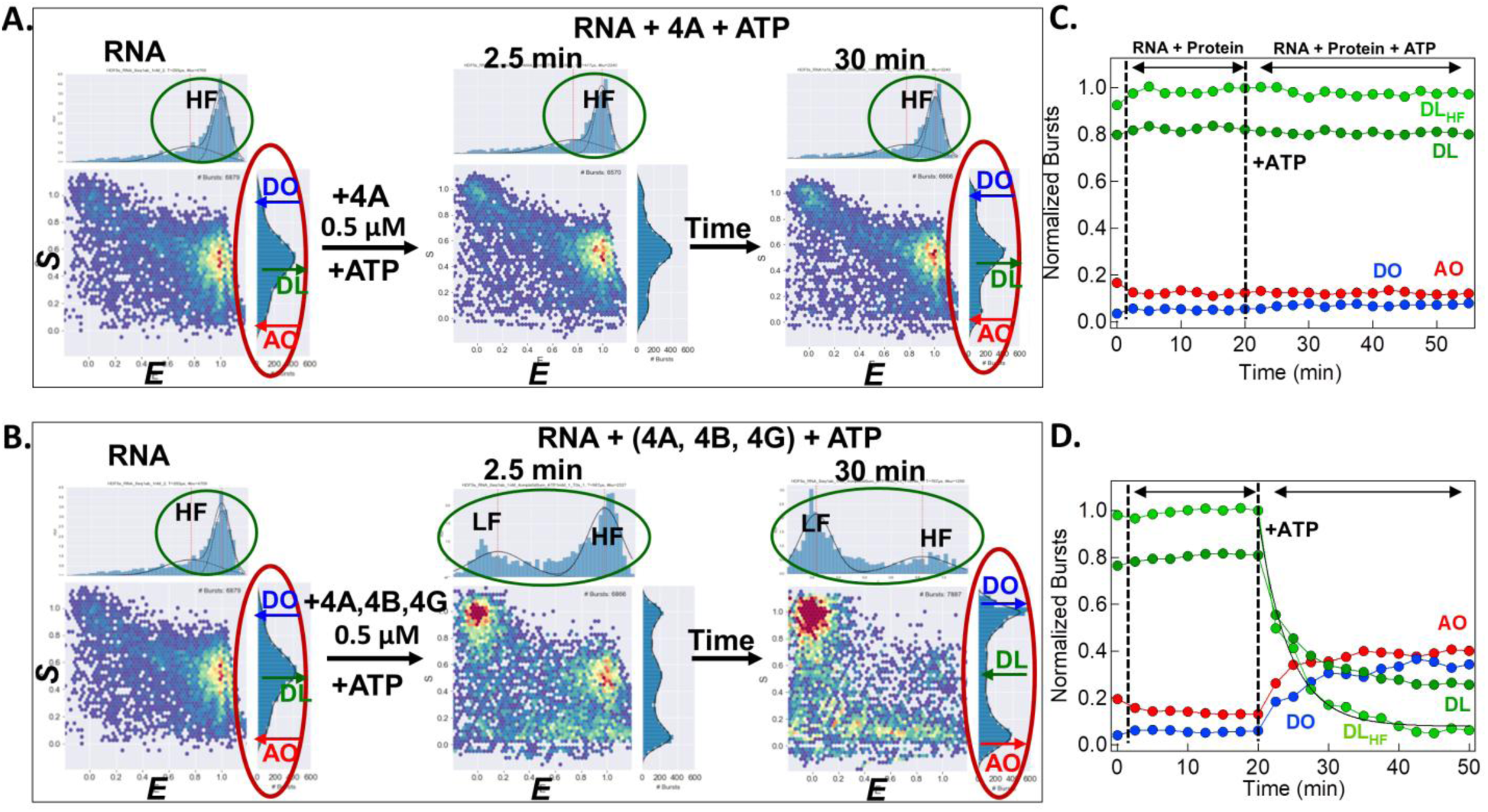
RPCs are functionally coupled and crucial to helicase activity. (A) 2D histogram of *S* vs. *E* showing donor-only (DO, Cy3, *S* = ∼1.0), doubly-labelled (DL, *S* = 0.2 – 0.8), and acceptor-only (AO, Alexa647, *S* = ∼0.0) population along with the FRET efficiencies (X-axis, *E,* green ellipse) and stoichiometry (Y-axis, *S*) histogram (red ellipse). The left scatter plot shows dsRNA only with high FRET efficiency (green ellipse), high DL particles and low DO and AO particles before adding eIF4A. The middle plot shows dsRNA after 0.5 μM eIF4A and 1 mM Mg-ATP addition at 2.5 min and 30 min (right scatter plot). No change in FRET is seen (green ellipse). The FRET efficiencies are calculated from DL particles defined as exhibiting *S* in the range 0.25-0.8. (B) 2D histogram of *S* vs. *E* of RNA as in left scatter plot: before adding eIF4A, eIF4B, eIF4G, middle: eIF4A, eIF4B, eIF4G right after addition of Mg-ATP at 2.5 min and 30 min (right scatter plot). Increase in DO and AO population and decrease in DL population (red spherical) as well as change in FRET efficiency from high FRET (HF) to low FRET (LF) (green spherical) shows the onset of helicase activity. (C) The fraction of doubly-labelled (DL; green), doubly-labelled HF(DL_HF_; light green), donor-only (DO; blue), and acceptor-only (AO; red) particles are plotted as a function of time for dsRNA alone, following the sequential addition of eIF4A (0.5 µM) and subsequently Mg-ATP (1 mM). The doubly-labeled dsRNA population remains unchanged after ATP addition, indicating no detectable helicase activity by eIF4A. First and second vertical line represents the time when protein and Mg-ATP were added, respectively. (D) The fraction of bursts populations (DL, AO, DO), doubly-labelled high FRET (DL_HF_; light green), for RNA alone, following the sequential addition of eIF4A, eIF4G, eIF4B (0.5 µM) and subsequently Mg-ATP (1 mM). Decrease in the doubly-labeled dsRNA population and corresponding increase in the DO and AO populations after ATP addition shows cofactor-stimulated helicase activity by eIF4A. Exponential is fit (black) to DL_HF_ to extract helicase rate and amplitude of DL unwound. First and second vertical line represents the time when protein and Mg-ATP were added, respectively.

The difference between the DL values (darker green data points in time course plots (e.g. Fig 2B) and the DL_HF_ values (lighter green dots) indicates the FRET changes within the RPCs due to partial unwinding of the dsRNA. This internal helicase rate *k*_f_ and its reversal, reannealing *k*_r_, were quantified by recurrent analysis of single particles (RASP) in which individual RPCs that diffused out of the confocal detection volume and then, by chance diffused back into the beam were compared during their first and subsequent fluorescence burst. The sum of *k*_r_ and *k*_f_ in RPCs is ∼70 s^−1^, *k*_r_ being ∼4-fold higher than *k*_f_. Toward the end of the recordings, at 30 min after ATP addition, the remaining DL particles had more balanced *k*_f_ and *k*_r_ values causing the partially unwound low-FRET component to grow relative to the fully double-stranded high-FRET component.

Upon ATP addition, the dsRNA-eIF4A-eIF4B-eIF4G complex shows both the formation of megaDalton RNA-protein clusters (Fig, 1D, 1F, 1H) and a marked enhancement of helicase activity relative to the dsRNA-eIF4A complex alone (Fig. 2D), suggesting a functional linkage between the RPC formation and the helicase activity. Motivated by this observation, we tested whether the clusters are also formed using the physiological eIF4F complex (eIF4A, eIF4B, full length eIF4G (eIF4G-FL) and cap binding protein eIF4E) in presence of dsRNA and we systematically examined the contribution of each individual component: eIF4A, eIF4B, full length eIF4G and eIF4E, to both cluster size and helicase activity, thereby testing quantitative correlation between RPC formation and strand separation (Fig. S2A–F).

eIF4A alone failed to induce either RNA-protein clustering or helicase activity upon ATP addition (Fig. S2A). Likewise, the eIF4A-eIF4G-FL complex showed no detectable strand separation or change in diffusion rate, indicating the absence of clusters (Fig. S2B). In contrast, addition of eIF4B and eIF4A to the dsRNA probe resulted in a detectable ATP-dependent decrease in the doubly-labeled (DL) population together with a pronounced reduction in diffusion rate relative to dsRNA alone, consistent with both cluster formation and minor separation (Fig. S2C). eIF4A with eIF4G-FL and eIF4E plus dsRNA led to a minor but reproducible ATP-dependent decrease in the DL population without a corresponding change in diffusion, indicating slow helicase activity in the absence of clustering (Fig. S2D). The combination of eIF4A with both, eIF4G-FL and eIF4B plus dsRNA, produced markedly accelerated helicase activity and substantially slower diffusion, consistent with the formation of large RPCs (Fig. S2E). While inclusion of eIF4E further stimulated eIF4A helicase activity within the eIF4A-eIF4B-eIF4G-FL complex, it did not measurably alter cluster size (Fig. S2F), thereby indicating that enhancement of the helicase activity is not rigidly correlated to RPC size. Quantitative analysis across all of the conditions revealed a strong correlation between amplitude of DL and DL_HF_ change (Fig. S3A) and helicase rate (Fig. S3B) with cluster size as indicated from the diffusion coefficients (Fig. S3C, S3D). Enhanced helicase activity generally, coincided with RPC formation, but not perfectly with RPC size. eIF4B, with extensive IDRs, is a key factor in in driving RPC formation, identifying it as a key multivalent component that nucleates cluster formation of the eukaryotic initiation factor 4 complex.

The cap binding protein, eIF4E enhances helicase activity through a mechanism that is apparently distinct from RPC size. The experiments carried out using eIF4A, eIF4B and a truncated functional core of eIF4G (eIF4G) lacking the eIF4E binding site or with the full length eIF4G (eIF4G-FL). Under all these conditions, we found that the RPC formation was coupled to helicase activity. Unless otherwise stated, all the further experiments described below were performed using eIF4A in combination with eIF4B and truncated functional core eIF4G lacking the eIF4E binding domain.

### eIF4B seeds the formation RNA-protein clusters

The predominantly intrinsically disordered structure of eIF4B, is well suited to mediating multivalent RNA-protein interactions and promoting higher-order assembly (refs). We tested the the concentration dependence of eIF4B activity in cluster formation and helicase activity, by varying eIF4B concentration while keeping eIF4A and eIF4G constant (0.5 µM each), together with 0.5 nM dsRNA and 1 mM Mg-ATP. (Fig. 3) Titration of eIF4B up to 500 nM revealed a progressive increase in RPC size (Fig. 3A-C), accompanied by gradual acceleration of both the amplitude and rate of helicase activity as indicated by decrease in the DL population (Fig. 3D-E)

**Figure 3.**
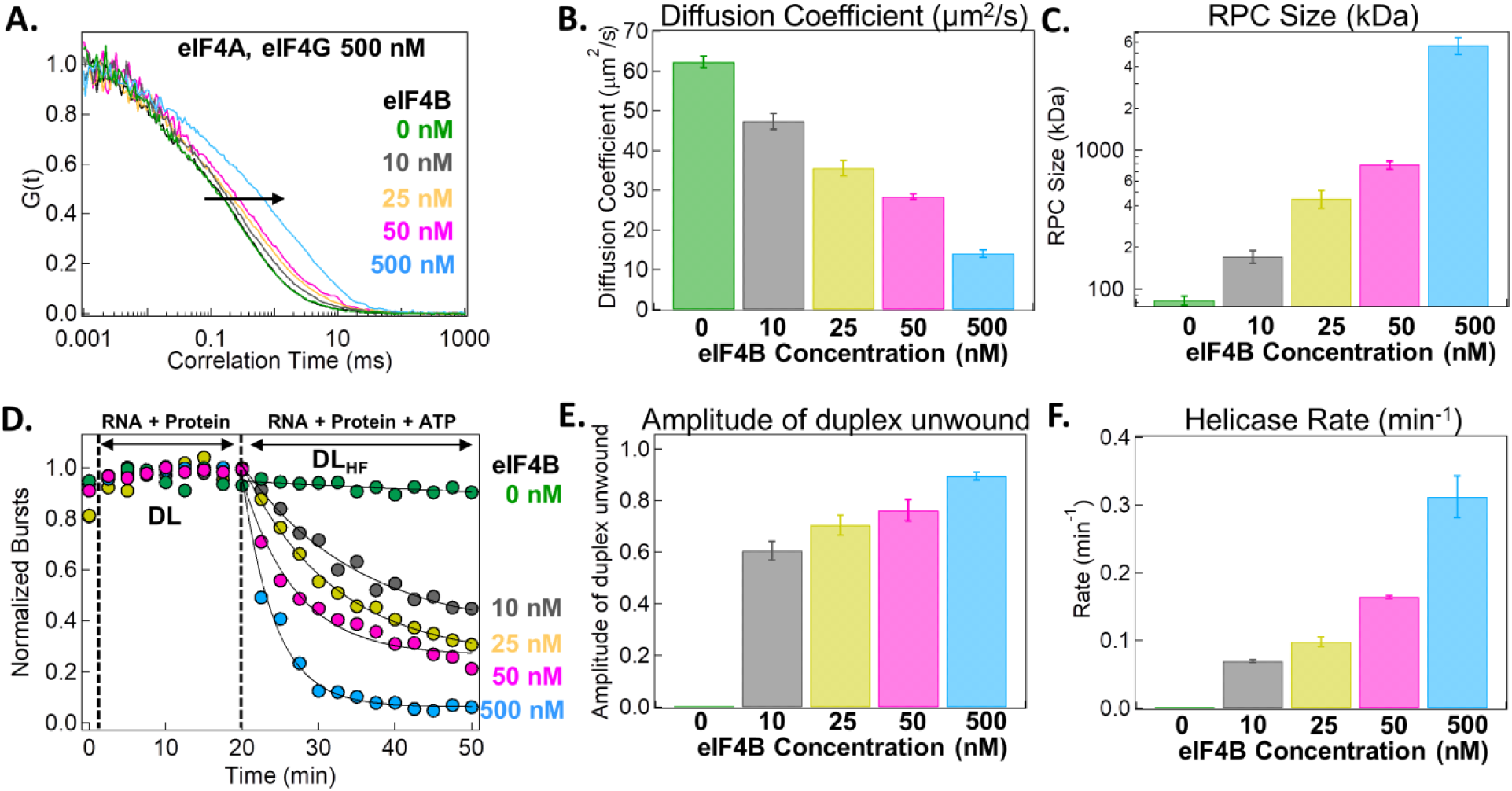
Titration of intrinsically disordered eIF4B reveals its role in nucleating RNA-protein clusters. (A) Normalized auto-correlation obtained from the FCS measurements are plotted for 0.5 nM dsRNA only (green), following the addition of eIF4A, eIF4G (0.5 µM each), and eIF4B 0 nM (dark green), 10 nM (grey), 25 nM (yellow), 50 nM (pink) and 500 nM (cyan) after 1 mM Mg-ATP addition. (B) Diffusion coefficient (µm^2^/s) extracted by fitting the auto-correlation curves following ATP addition to the dsRNA-eIF4A-eIF4G and subsequent titration of eIF4B at increasing concentrations. (C) RNA-protein cluster size calculated from the obtained diffusion coefficients. Increasing concentration of eIF4B induces larger cluster formation. (D) Amplitude of doubly-labeled (DL_HF_) population change following ATP addition to the dsRNA-eIF4A-eIF4G and subsequent titration of eIF4B at increasing concentrations (0, 10, 25, 50, 500 nM). (E) Amplitude of duplex unwound and (F) rate of the helicase activity following ATP addition to the dsRNA-eIF4A-eIF4G complex and subsequent titration of eIF4B at increasing concentrations.

To test the possibility that eIF4B undergoes self-aggregation driven by its IDRs in the sub-micomolar concentration range, we performed control diffusion measurements with dsRNA added to eIF4B alone. Incubation of eIF4B (0.5 or 1 µM) with 0.5 nM dsRNA produced no detectable change in diffusion relative to dsRNA alone (Fig. S4A), indicating the absence of eIF4B-driven aggregation and RPC formation under these conditions. Similarly, diffusion measurements of 1 nM Cy3-labeled eIF4B (Cy3-eIF4B) in the presence of excess unlabeled eIF4B (1 µM) showed no change compared with Cy3-eIF4B alone (Fig. S4B), ruling out the self-association of eIF4B at this concentration.

eIF4H is a paralog of eIF4B that retains similar structured RNA recognition motifs (RRMs) but contains much smaller intrinsically disordered regions relative to eIF4B (Fig. S5A, S5D). To gauge the influence of the disordered regions, we tested eIF4A, eIF4G, and eIF4H (0.5 µM each) with 0.5 nM dsRNA in the presence of 1 mM Mg-ATP (Fig. S5B) and quantified donor-only (DO), doubly-labeled (DL), and acceptor-only (AO) populations as the function of time (Fig. S5C). Under these conditions, eIF4H showed markedly slower strand separation rate compared with eIF4B at equivalent concentrations (Fig. S5E, S5F). eIF4H at 0.5 and 1 µM in the presence of eIF4A and eIF4G-core (0.5 µM each) formed smaller concentration-dependent RPCs (Fig. S6C, S6D) that correlated with lower helicase activity than that stimulated by eIF4B (Fig. S6A, S6B, S6D, S6E). Finally, control experiments with eIF4H (0.5 µM) alone showed no effect on diffusion rate of dsRNA (Fig. S7A). While eIF4H promotes helicase activity and smaller clusters, the broader IDRs of eIF4B are required for robust cluster nucleation and maximal helicase enhancement.

### RNA-eIF4B interaction critical for cluster formation and helicase activity

To assess the contribution of RNA-eIF4B interactions to helicase activity and RPC formation, we created a variant eIF4B with a targeted point mutation (F139A) within the structured RNA-recognition motif (RRM) which is expected to reduce the RNA binding affinity (Fig. 4 A). We measured diffusion of clusters containing eIF4A, eIF4G, and either wild-type or mutant eIF4B (0.5 µM each) in the presence of 0.5 nM dsRNA and 1 mM Mg-ATP. Complexes formed with mutant eIF4B exhibited much faster diffusion, indicating reduced cluster size, compared with those containing wild-type eIF4B (Fig. 4B, 4C). Consistent with this observation, the rate constant for helicase induced decrease in the DL population markedly lower for the mutant-eIF4B complex relative to the wild-type complex (Fig. 4E, 4F). Mass photometry measurements independently confirmed a difference in assembly size, with molecular mass estimates closely matching those inferred from FCS (Fig. 4D). Finally, transmission electron microscopy of negatively stained samples containing eIF4A, eIF4G, and mutant eIF4B (0.5 µM each) in the presence of dsRNA and Mg-ATP revealed the formation of distinct, round, homogeneous clusters that were markedly smaller than those formed with wild-type eIF4B (Fig. 4G). Thus, the affinity of the RRM-mediated interaction between eIF4B and RNA is critical for robust cluster formation and efficient helicase activity, again revealing a functional coupling between clustering and helicase activity.

**Figure 4.**
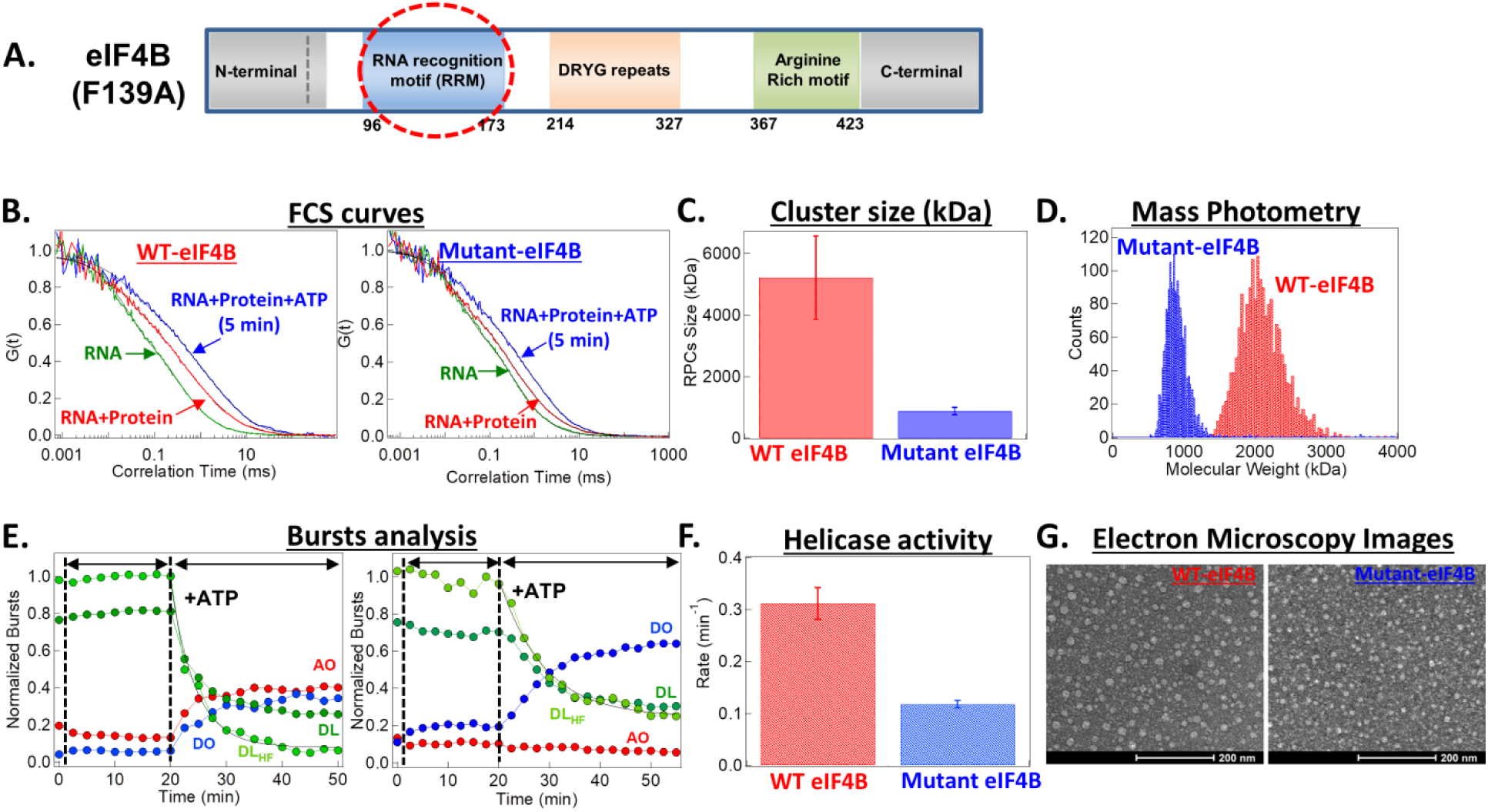
RNA recognition motifs (RRM) of eIF4B is critical for cluster formation and helicase activity. (A) Schematic representation of wild-type eIF4B domain architecture, with a point mutation (F139A) introduced in the structured RNA recognition motif (RRM) to generate mutant eIF4B. (B) Auto-correlation curve obtained from the FCS measurement for 0.5 nM dsRNA only (green), after addition of eIF4A, eIF4G, WT eIF4B or mutant eIF4B (0.5 μM each) (red) following subsequent addition of 1mM Mg-ATP (blue). WT eIF4B complex diffuses slower than mutant eIF4B complex indicating the difference in the cluster size. (C) Cluster size in kDa for dsRNA-eIF4A-eIF4B(WT)-eIF4G complex (red) and dsRNA-eIF4A-eIF4B(mutant)-eIF4G complex (blue). Data are presented as the mean ±SD of 6 independent repeats. (D) Histograms of apparent molecular weights (kDa) of 0.5 μM eIF4A-eIF4B(WT)-eIF4G (red) and 0.5 μM eIF4A-eIF4B(mutant)-eIF4G (blue) with 0.5 nM dsRNA were obtained using mass photometry. (E) The fraction of doubly-labelled (DL; green), doubly-labelled HF (DL_HF_; light green), donor-only (DO; blue), and acceptor-only (AO; red) particles are plotted as a function of time for dsRNA alone, following the sequential addition of eIF4A, eIF4G, WT eIF4B or mutant eIF4B (0.5 µM each) and subsequently Mg-ATP (1 mM). Decrease in the double-labeled dsRNA population and subsequent increase in the DO and AO population after ATP addition shows cofactors stimulated helicase activity by eIF4A. Exponential is fit (black) to DL_HF_ to extract helicase rate and amplitude of DL unwound. First and second vertical line represents the time when protein and Mg-ATP were added, respectively. (F) Helicase rate for dsRNA-eIF4A-eIF4B(WT)-eIF4G complex (red) and RNA-eIF4A-eIF4B(mutant)-eIF4G complex (blue) obtained by fitting the change in DL_HF_ using an exponential equation. (G) Representative electron microscopy image of negatively stained 0.5 μM eIF4A-eIF4B(WT)-eIF4G (left panel) and 0.5 μM eIF4A-eIF4B(mutant)-eIF4BG with 0.5 nM dsRNA (right panel). Scale bar, 200 nm.

RASP analysis showed partial helicase and reannealing of the RNA within the RPCs containing mutant eIF4B similar to that of wild type RPCs. Also similar, the helicase rate, *k*_f_, was ∼4-fold higher than the reannealing rate promptly after ATP addition, but the two rate constant values were more balanced in the DL particles still remaining 30 min later. The number of mutant DL particles remaining at 30 min was greater, however, than WT RPCs due to the reduction of the overall helicase activity of the mutant (Fig. 4F).

### Diffusion rate of WT- and mutant-eIF4B in live cells reflects RRM governed clustering

To assess whether eIF4B diffuses freely or is incorporated into RPCs in the cellular environment and whether the differences in cluster size observed between wild-type (WT) and mutant eIF4B *in vitro* also exist in cells, we expressed eGFP-tagged WT-eIF4B, eGFP-tagged mutant-eIF4B, and eGFP alone at comparable levels in HeLa cells (Fig. 5A). Similar low levels of expression for all three transformations were obtained under basal conditions in the absence of tetracycline induction, due to leakage of repression at the Tet inducer (Fig. 5A). Cells were maintained in an environmental chamber at 37 °C and 5% CO₂. Fluorescence correlation spectroscopy revealed that WT eIF4B exhibited the slowest intracellular diffusion, particularly at longer timescales (Fig. 5B), with an apparent diffusion coefficient of 0.63 ±0.03 µm² s⁻¹ (mean ± s.e.m., n = 23). Mutant eIF4B diffused substantially faster than WT eIF4B, with an apparent diffusion coefficient of 1.01 ±0.07 µm² s⁻¹ (mean ± s.e.m., n = 23), while GFP alone displayed the fastest diffusion (22.2 ±0.81 µm² s⁻¹, (mean ± s.e.m., n = 26)) (Fig. 5C). Diffusion coefficient for GFP *in vitro*^30^ is ∼80 µm² s⁻¹ approximately 4-fold faster than within the cells^31^. Diffusion of WT eIF4B is approximately 10-fold slower in the cells than *in vitro*, indicating that it diffuses in a large complex with other cellular components, presumably the other initiation factors, similar to the *in vitro* RPCs. Faster diffusion of the mutant eIF4B indicates that it’s cluster size is smaller than WT’s, similar to what is observed *in vitro*, or that fewer mutant eIF4B molecules enter clusters. In either or both cases the faster diffusion of the mutant with weaker RNA binding affinity indicates that interaction with RNA is important in formation of the diffusing particles as it is *in vitro*. Together, these results strongly suggest that WT-eIF4B enters slowly diffusing RPCs in cells containing RNA, consistent with its *in vitro* behavior. The intracellular assemblies are likely heterogeneous and may incorporate additional cellular components, for instance ribosomal subunits.

**Figure 5.**
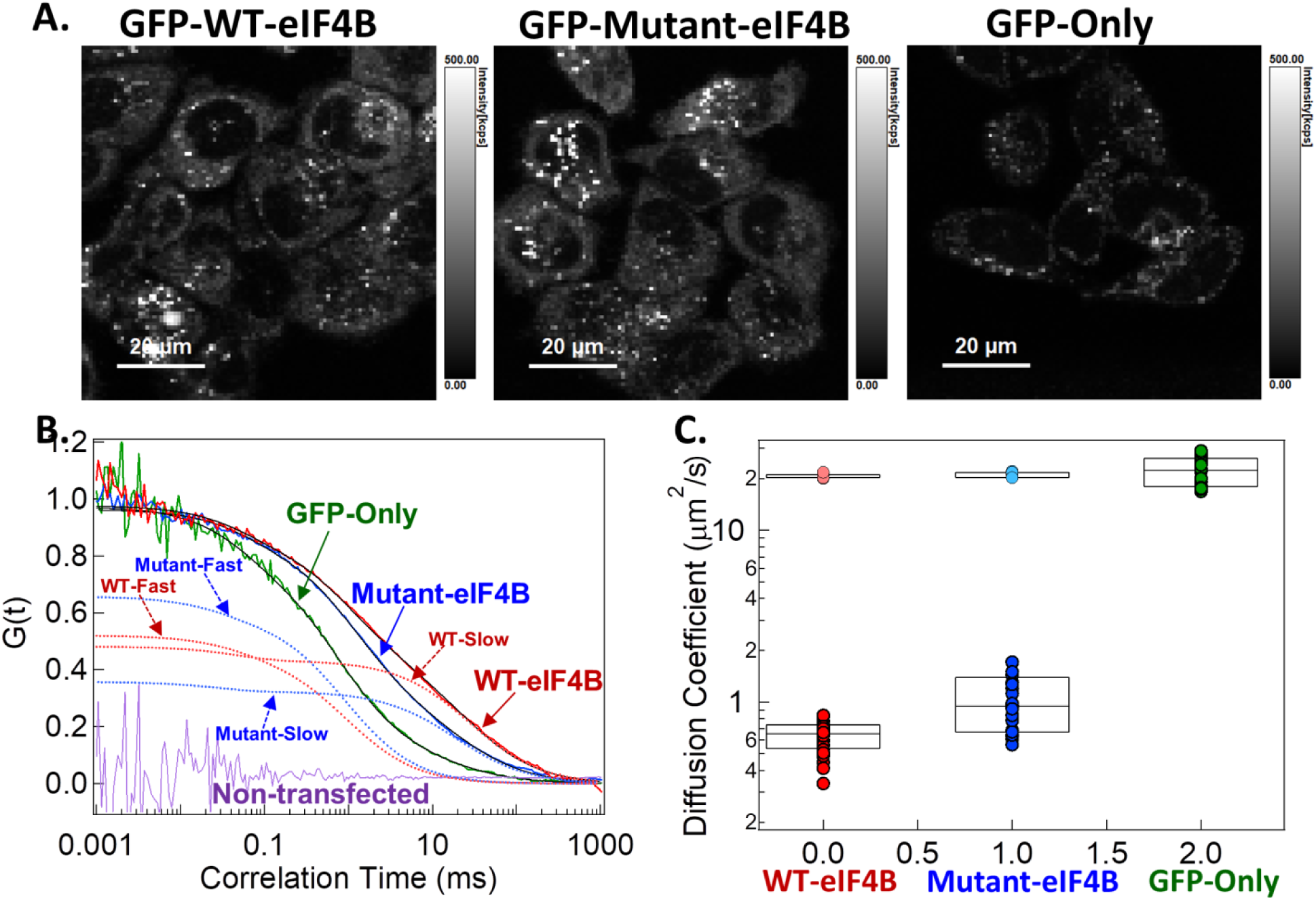
Distinct cellular diffusion states of wild-type and RRM-mutant eIF4B reveal RNA recognition motif-dependent cluster formation in HeLa cells. (A) Fluorescent intensity-based images of HeLa cells expressing GFP tagged WT eIF4B, GFP tagged mutant eIF4B and GFP-only. The grey bar on the right side shows fluorescent intensity in kilo counts per second (kcps). Scalebar, 20 µm. (B) Normalized auto-correlation curves of GFP-only (green), GFP-tagged mutant-eIF4B (blue) and GFP-tagged WT-eIF4B (red). The fit to the data is shown in black. Non transfected parent cell lines were used as control showing zero correlation (purple). Both WT eIF4B and mutant eIF4B correlation curves show a distinct fast and slow components. Notably, the mutant eIF4B (blue dashed line) exhibits a higher amplitude of its faster component but a lower amplitude of the slower component (blue dashed line) compared to the WT eIF4B (red dashed line). (C) Apparent diffusion coefficients (D_app_) (µm^2^/s) of GFP-tagged WT eIF4B (red), GFP-tagged mutant eIF4B (blue) and GFP-only (green). Mean values were calculated from n = 42. The correlation curves are fit using two diffusing species and triplet. The fast diffusing species in WT eIF4B (coral) and mutant eIF4B (cyan) diffuses with a similar rate to that of GFP-only construct.

## Discussion

The mechanical unwinding of structured 5’ untranslated regions (UTRs) by eIF4A, a highly conserved DEAD-box helicase, is a crucial step in eukaryotic translation initiation. While generally considered an isolated, non-processive monomer that relies on stochastic collisions to resolve RNA secondary structure sequentially^10, 32^, our current findings offer a new perspective that expands upon this classic view. Using a multi-level biophysical approach that combines quantitative single molecule fluorescence resonance energy transfer (smFRET), fluorescence correlation spectroscopy (FCS), mass photometry (MP), and negative-stained electron microscopy (EM), this work shows that eIF4A, functions as nanometer-scale RNA-protein clusters (RPCs) in presence of the two accessory proteins, eIF4G and eIF4B. These assemblies, formed within the physiological concentration range, suggest that eIF4A mediated catalysis is not stochastic, but rather an organized process functionally coupled to the formation of these native clusters. The markedly enhanced RNA unwinding rate within the RPC helps explain how eIF4A transitions from a low-turnover state, on its own, to a highly active helicase in presence of the accessory proteins.

Earlier studies have shown that isolated eIF4A is weak and non-processive helicase^2, 33^. Without the accessory factors, eIF4A on its own destabilizes a only 1 to 3 base pairs per binding cycle before dissociating^10^. This low activity demonstrates that its catalytic cycle is not coupled tightly to unwinding and that ATP hydrolysis fails to drive sustained strand separation unless modulated by scaffolding partners eIF4G and eIF4B^34^. To overcome this weak activity *in vitro*, previous single molecule studies relied on high stoichiometric excess, utilizing micromolar concentrations of protein against picomolar RNA concentrations to observe detectable and accelerated ATP-dependent unwinding in the presence of the cofactors^10, 34^. Combining a short dsRNA having a ssRNA overhang at nM concentration with eIF4A, eIF4B, eIF4G and ATP in the μM range increased unwinding activity by several folds. Rather than the high concentration used in previous studies being just an experimental requirement to drive a weak enzyme, the current work provides the dynamic rationale for the concentration dependence by demonstrating that the functional, high activity of eIF4A with its cofactors is natively achieved through regulated organization into nanometer-scale RNA-protein clusters.

In the presence of RNA and ATP but without eIF4B and eIF4G, mass photometry data showed that eIF4A forms discrete high molecular weight species ∼360 kDa (Fig. S1C). Given eIF4A’s molecular mass of ∼46 kDa, this complex corresponds to a multimeric oligomer. This observation aligns with Schmidt et al.^35^, which showed that eIF4A undergoes sequence- and ATP-dependent multimerization on single-stranded polypurine motifs at physiological concentrations. While our mass photometry data resolved distinct multimers, the FCS results monitoring RNA Alexa647 autocorrelation in the absence of the cofactors did not show a corresponding suppression of RNA’s diffusion coefficient (Figure 1C). This difference could point to a highly dynamic or transient assembly in which eIF4A, loosely packed on the flexible ssRNA does not significantly increase its overall hydrodynamic radius. Crucially, however, our data shows that these multimers remain in catalytically suppressed state incapable of unwinding RNA until reorganized by the accessory factors (Figure 2C).

Introducing eIF4B and eIF4G triggers a dramatic structural reorganization. The addition drives a substantial hydrodynamic shift in FCS (Fig. 1D, 1G) and yields distinct ∼875 kDa assemblies detected by mass photometry (Fig. S1C). Upon subsequent addition of Mg-ATP, the components assemble into RNA-protein clusters with ∼2.1 MDa by mass photometry (Fig. S1C), which aligns well with a corresponding large reduction in RNA diffusion rate indicative of higher molecular weight (Fig. 1G) by FCS. Strikingly, this stepwise growth directly scales with the cooperative activation of the helicase where eIF4A is systematically transitioned from weak strand separation to a highly regulated, organized and accelerated catalytic helicase. To decouple the individual contributions of the initiation factors to formation of RPCs and the accelerated activity, we analyzed sub-complex assemblies. Interestingly, addition of the scaffold, eIF4G, to eIF4A plus RNA, after ATP addition, yielded a mass similar to eIF4A-RNA-ATP using mass photometry in line with the FCS results (Fig. S1C, S2B). Similarly, mass photometry and FCS of eIF4B alone, evaluated either in absence or presence of dsRNA, yielded a consistent mass of ∼152 kDa (Fig. S1C). It was exclusively when all three initiation factors, eIF4A, eIF4G and eIF4B were combined, we observed an ATP dependent transition into ∼2.5 MDa RPCs.

Our understanding of RNA-eIF4A-eIF4G-eIF4B RPCs, well below the concentration (*C*_sat_) leading to liquid-liquid phase separation and μm-scale liquid droplets suggests that the multi-molecular assemblies exist in a pre-percolated state. In this sub-binodal regime, molecular networking occurs in a dual-transition process termed phase separation coupled to percolation (PSCP)^16, 17^. The hypothesis is that these assemblies rely on a “stickers-and-spacers” architecture, where folded domains and short motifs act as associative elements, while disordered regions serve as flexible spacers that modulate network connectivity and viscoelasticity^36^. Biophysical and structural characterization of multi-valent interactions over a broad class of RNA-binding proteins including other DEAD-box helicases^22^, DDX4^23^, Ded1p^25^, DDX6^25^, the FET family (FUS^20^, EWS/TAF15^21^), and critical scaffolds (G3BP1^13^, TDP-43^37^, FMRP^38^, NPM1^39^) collectively demonstrate that underlying molecular networks exist across an operational continuum, depending on the percolation (*C*_perc_) and phase separation (*C*_sat_) thresholds, rather than in a binary transition into macroscopic droplets^40^. Ultimately, the nanometer-scale clusters serve as catalytically optimized, high-density hubs that accelerate enzymatic turnover while avoiding the risk of kinetic arrest and exclusion from substrates associated with μm-scale macroscopic condensates.

Our data shows that the predominantly disordered eIF4B serves as the critical component driving both cluster size and catalytic acceleration (Fig. 3A-F) presumably due to its largely disordered structure. The crucial involvement of disordered segments of eIF4B interacting with the eIF4A-eIF4G core to drive cluster formation is similar to the mechanism of DEAD-box helicase DDX3X/Y RPC stimulated clustering and dsRNA unwinding^22^. Swain et al.^27^ reported that human eIF4B undergoes self-association *via* its C-terminal IDR in the concentration range between 1 and 25 μM, whereas our results at lower physiological eIF4B concentrations require eIF4B binding to the eIF4A-eIF4G complex to obtain RPCs (Fig. S2A-F).

Within the stickers and spacers paradigm, while eIF4G often acts as the primary multivalent scaffold, the closed conformation state of eIF4A drastically increases its affinity for RNA, making it a strong dynamic sticker. Comparison of eIF4B with its human paralog, eIF4H (Fig. S5A-F) identified other associative (sticker) regions. Although eIF4B and eIF4H share structural homology within their core domains, eIF4H lacks extensive disordered sequences, including the self-associating DRYG domain and the C-terminal RNA-binding domain. eIF4H formed smaller RPCs with lower helicase activity than eIF4B (Fig. S6A-F). The size reduction was more than could be explained by the lower molecular weight of eIF4H (∼25 kDa) compared to eIF4B (∼77kDa), thus suggesting that the C-terminal RRM and the DRYG sequences are significant. The RNA recognition motif (RRM) of eIF4B is another associative component as demonstrated by our result that mutating eIF4B phenylalanine at 139^th^ position to alanine reduced RPC size and helicase activity (Fig. 4B-G)

The apparent size difference observed between wild-type and mutant eIF4B RPCs in vitro also persists in HeLa cells, further supporting a critical role for the eIF4B RRM in regulating higher-order assembly (Figure 5A–C). The diffusion coefficients of GFP-tagged WT and mutant eIF4B in HeLa cells are significantly slower than those measured in vitro, even after correcting for higher intracellular viscosity, suggesting interactions with additional cellular components. Together, these findings support the cellular relevance of eIF4B-dependent clustering and raise the possibility that related higher-order assemblies may contribute to broader translational RNP organization in cells.

Full unwinding and separation of RNA strands into separate clusters was monitored in our study by a decrease in the doubly-labeled (DL) population of particles, accompanied by a simultaneous increase in the donor-only (DO) and acceptor-only (AO) populations as the strands split. In the presence of ATP, remaining DL clusters exhibit high and low FRET efficiency components which indicate partial unwinding. Burst Variance Analysis (BVA) showed rapid exchange between the two DL particle FRET states in RPCs formed from mutant eIF4B indicatting partial dsRNA unwinding and rapid reannealing within the RPC. But even though the overall rate of full helicase activity is higher with WT eIF4B, surprisingly, BVA did not indicate substantial FRET state exchange in WT eIF4B RPCs. We reasoned that partial unwinding and reannealing might be too slow to detect by BVA which is limited to exchange processes on the time scale of particle occupancy in the PicoQuant detection volume (0.5 – 2 ms). A powerful way to extend the time scale for estimating exchange rates between FRET states is recurrence analysis of single particles (RASP) which considers particles that diffuse out of the detection volume and then, by chance, re-enter it. This analysis also sheds light on whether individual RNA molecules undergo conformational fluctuations, transiently shifting between fully and partially wound states. While mutant eIF4B showed a slower unwinding rate and a reduced overall transition from a high to a low FRET state, these delayed transitions suggest that RPCs containing the mutant protein are less efficient. The mutant eIF4B may likely require more ATP cycles to fully unwind a given length of dsRNA compared to WT eIF4B.

In conclusion, our study expands the working framework of eIF4A from an isolated helicase to a regulated and organized RNA–protein cluster state. These nanometer-scale assemblies link higher-order molecular organization to increased catalytic output by creating a local biochemical environment favorable for repeated partial RNA unwinding and efficient strand separation. More broadly, these findings provide a framework for understanding how non-processive DEAD-box helicases may achieve efficient RNA remodeling in the crowded cellular environment.

## Acknowledgements

We thank Sophie Kiss for obtaining negative stained TEM images.

## Author contributions

M.S. and N. V. performed the molecular cloning and protein purifications. H.S. performed the time-resolved spectroscopy experiments and analyzed the data. All authors participated in the conception of ideas, planning of experiments, discussion of results, and writing/editing the manuscript.

## Data and material availability

Data are available in the manuscript or the supplementary material. Materials are available on request to the corresponding author.

## Methods

### Protein expression and purification

Recombinant human eIF4A^2^, eIF4H^2^, eIF4G-core^2^, eIF4E^28^ and eIF4G-FL (165-1599) proteins were expressed and purified as previously described. The gene encoding human eIF4B was commercially synthesized (GenScript) and subcloned into the parent vector via NdeI and XhoI restriction sites of pET28c vector. The single native cysteine residue was then mutated to serine (C457S) and a new cysteine was introduced (S183C) using site-directed mutagenesis. F139A mutation in eIF4B RNA recognition motif (RRM) was then further introduced. Resulting N-terminal His-tagged eIF4B (WT or mutant) was expressed in E. Coli BL21 (DE3), and purified as essentially described previously for eIF4B expressed in insect cells^28^.

### Plasmid construction and generating inducible cell lines

To generate stable mammalian cell lines, the coding sequence of WT human eIF4B containing an N-terminal EGFP tag was subcloned into pcDNA5/FRT/TO expression vector (Invitrogen) utilizing BamHI and XhoI restriction sites. The EGFP-tagged eIF4B mutant (F139A) plasmid was subsequently generated by inserting a mutant RRM fragment, synthesized by GenScript, into the parent EGFP-WT vector via CIaI and BstBI restriction sites. All primers were purchased from Integrated DNA Technologies (IDT), and the plasmid sequences were verified by Sanger sequencing (Azenta Life Sciences and Quintara Biosciences).

### Cell culture and maintenance

HeLa Flp-In™ T-Rex cell lines (a generous gift from Drs. Elena Dobrikova and Matthias Gromeier, Duke University Medical Center) were maintained in Dulbecco’s Modified Eagle Medium (DMEM) supplemented with 10% fetal bovine serum (FBS) and cultured at 37 C in a humified atmosphere with 5% CO_2_. For the in cell FCS measurements, the standard growth medium was replaced with Gibco FluoroBrite DMEM supplemented with 2% FBS for at least 1 hour prior to data acquisition to minimize background fluorescence.

### Transfection

Stable cell lines expressing EGFP-tagged eIF4B WT or the eIF4B mutant were generated using the Flp-In™ system. Transfections were performed in 6-well plates using Opti-MEM Reduced Serum Medium (Thermo Fisher Scientific) supplemented with 3% FBS. For each transfection, 1.5–3.0 µg of the appropriate pcDNA5/FRT/TO plasmid and 1 µg of the pOG44 recombinase expression plasmid were diluted in 100 µL of Opti-MEM. Concurrently, 6 µL of X-tremeGENE 9 DNA Transfection Reagent (Roche) was diluted in 100 µL of Opti-MEM and incubated for 5 minutes at room temperature. The DNA and lipid dilutions were then combined and incubated for 30 minutes at room temperature to allow complex formation. The resulting DNA-lipid mixture was added dropwise to the cells and incubated for 5 hours. Following transfection, stable integrants were selected using Blasticidin and Hygromycin selection in accordance with the manufacturer’s guidelines to establish the final inducible lines.

### Multi-parameter confocal time resolved spectroscopy

The single molecule measurements utilizing fluorescent probes were performed on multi-parameter confocal fluorescence time resolved spectroscopy (MicroTime-200; PicoQuant, GmbH) based on the principle of time correlated single photon counting (TCSPC) within ∼femtoliter (fL) observation volume. All the measurements were done in translation buffer (20 mM Hepes-K pH 7.5, 45 mM KCl, 90 mM K acetate, 2.2 mM Mg acetate, 0.1 mM spermidine, 1 mM DTT). A duplex RNA labeled with Cy3 and Alexa647 were used for all the measurements, where a 24-mer (5’-Cy3/GUUUUUUAAUUUUUUAAUUUUUUC/-3’) was annealed with 44-mer (5’-/GAACAACAACAACAACAACAGAAAAAAUUAAAAAAUUAAAAAAC/Alexa647/-3’) (IDT). For the fluorescence measurements, the Cy3 (donor) and Alexa647 (acceptor) dyes on the labeled RNA were excited using pulsed diode lasers (PicoQuant LDH-D-TA-532 and LDH-D-TA-637) operating at 20 MHz in pulsed-interleaved excitation (PIE) mode. The excitation beams were focused 20 μm above the coverslip interface using a ZT532/637 dichroic filter (Chroma Technology) and an Olympus UPLanSApo 60x/1.2 water immersion objective, with laser powers maintained below 20 μW. Emitted fluorescence passed through a 50 μm pinhole and was separated into vertical and horizontal polarization paths via a polarizing beam splitter. Within each path, T635 lpxr dichroic filters (Chroma Technology) split the signals into Cy3 and Alexa647 channels, which were further isolated using ET582/64 and ET690/70 bandpass filters, respectively. The resulting polarized, wavelength-selected photons were detected by four single-photon avalanche photodiode (SPAD) detectors and recorded using a HydraHarp TCSPC time-interval analyzer (PicoQuant).

Data was colllabeled RNA alone, followed by multiple 2.5 minute acquisitions over 30 minutes’ post-protein addition, and ∼40 minutes recording after adding of 1 mM Mg-ATP. These *in vitro* experiments were conducted in Nunc Lab-Tek chambers (ThermoFisher-155411) featuring borosilicate coverslip bottoms, modified with a circular punched silastic sheets to reduce the chamber’s volume to 40 uL, which were then passivated for a minimum of 4 hours at room temperature with a 50% (w/v) PEG-8000 solution and rinsed 3-4 times with nuclease free water followed by 2 washes with translation buffer. Confocal detection volumes for the 532 nm and 637 nm lasers were calibrated using Rhodamine6-G and atto637N dye standards.

### Single molecule PIE FRET

The raw PIE datasets collected on PicoQuant were first converted into structured open-source photon HDF5 format. This conversion was done using the python-based ‘phconvert’ module within the ‘FRETBursts’ package. Because the FRET calculations do not require polarization specificity, the photon counts recorded across both the vertical and horizontal detection paths were merged for both the donor and acceptor channels. The background rate was computed across consecutive 30 seconds intervals to account for temporal fluctuations. Background corrected photons were then subjected to multiple corrections, accounting for spectral crosstalk (leakage), direct excitation of acceptor by donor laser (direct excitation), differences in localized laser power, and the dye absorptivity, quantum yields, and relative detector sensitivities. By maintaining the labeled duplex RNA concentration to ∼0.5 nM, we limited the average molecular occupancy within the observation volume to 0.2-0.3 molecules at any given moment. To reliably isolate individual freely diffusing particles from the baseline noise, we used a sliding window algorithm. A burst was identified when a sequence of *m* = 8 consecutive photons arrived at a rate exceeding the background threshold by a factor of five (*F* = 5), provided the total intensity of the event contained at least *L* = 15 photons.

Following all necessary background and instrumental corrections, each single-molecule burst yields three distinct photon streams: *F_Dem/Dex_*(Donor emission resulting from donor excitation), *F_Aem/Dex_* (Acceptor emission resulting from donor excitation), *F_Aem/Aex_* (Acceptor emission resulting from direct acceptor excitation). Using these corrected photon counts, the apparent FRET efficiency (*E*) and stoichiometry (*S*) are calculated for each individual burst using Equations 1 and 2:

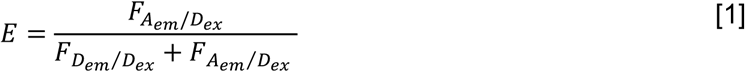

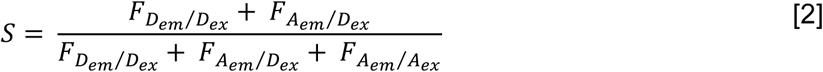

Individual photon bursts generated by particles diffusing through the detection volume were analyzed using the FRETBursts framework, with the pulse-interleaved excitation (PIE) scheme of the confocal spectrometer to classify each burst according to the relative contributions of donor (Cy3) and acceptor (A647) fluorescence. This analysis enables assignment of each detected particle as donor-only (DO), acceptor-only (AO), or doubly-labeled (DL) based on its fluorescence stoichiometry (S). In this representation, DO species (short strand labeled with Cy3) exhibits S ≈ 1, AO species (long strand labeled with Alexa647) appear at S ≈ 0, and DL duplex particles containing equal numbers of donor and acceptor probes appear at S ≈ 0.5. For the DL population, FRETBursts additionally computes the FRET efficiency (E), which reports on the distance between the Cy3 and Alexa647 fluorophores and thus serves as a molecular ruler for the dsRNA conformation. These assignments for ∼6000 particles detected within 2.5 min are displayed as two-dimensional histograms of stoichiometry (S, vertical axis) versus FRET efficiency (E, horizontal axis) in Fig. 2A and B. The one-dimensional histograms to the right of each panel show the relative abundance of DO, DL, and AO particles, while the histogram above each plot shows the FRET efficiency distributions specifically for the DL population (S ≈ 0.5). For clarity in the main text figures, these three particle populations are expressed and normalized using Equation 3.

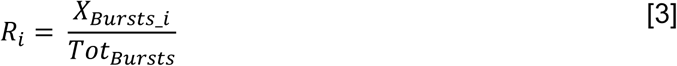

Where, *R_i_* is the normalized DO or AO or DL population, *X_Bursts_i_* is the DO or AO or 2*DL population after burst categorization, *Tot_Bursts_* is the total number of bursts in the *E vs. S* histogram, given as DO + 2*DL + AO.

The DL particles were categorized into high-FRET (HF) and low-FRET (LF) components using a FRET efficiency threshold of 0.75. Following the addition of ATP, dsRNA strand separation causes the FRET efficiency peak to shift towards zero. Consequently, the DO population tail extends into DL population region at stoichiometry values below S = 0.8, introducing a population artifact. To correct for this contamination and accurately capture the kinetics of the DL population, the time-dependent change in HF fraction was normalized by the total baseline DL population before ATP addition using Equation 4:

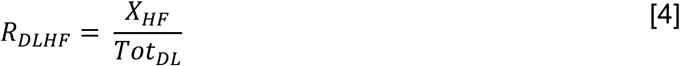

Where, *X_HF_*is the occupancy of the high-FRET group within the FRET distribution, *Tot_DL_*is the total DL population prior to ATP addition.

### Fluorescence Correlation Spectroscopy (FCS)

Due to the close proximity of the Cy3 and Alexa647 dyes on the dsRNA, the FRET efficiency approached nearly 100 %. Consequently, the lack of detectable donor signal prevented the auto-correlation of donor signal as well as the cross-correlation analysis of donor and acceptor signal. Therefore, acceptor signal was auto-correlated to evaluate the bound state of dsRNA-protein complex both before and after ATP addition. Utilizing the acceptor fluorescence signals separated by a polarizing beam splitter, directed onto two SPADs, to auto-correlate eliminates the effect of uncorrelated detector after-pulses. Auto-correlation curves were fitted with a diffusion model containing one or two diffusing species including the triplet state (Equation 5):

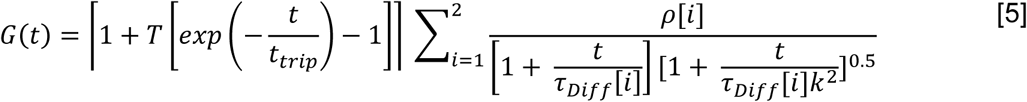

Where, *i* denotes the faster or slower diffusing species, represents the fractional contribution of the *i*^th^ diffusing species, *k* is structural parameter defined as the length to diameter ratio of the focal volume, 𝜏_𝐷_ indicates the diffusion time of the *i*^th^ diffusing species, *T* and 𝑡 _𝑟_ represent the triplet state fraction and the triplet relaxation time of the fluorophore. Diffusion time, obtained as correlation time at half-maximum amplitude (𝜏_1/2_) of the triplet-corrected auto-correlation curve, was used to calculate the apparent diffusion coefficient (*D_app_*) using the relation, *D_app_* = *w*_0_ ^2^/4 𝜏_1/2_, where *w*_0_ represents the radius of the detection volume at the beam waist.

### In cell FCS

Stable HeLa cell lines expressing EGFP-eIF4B (WT or mutant) were harvested and seeded into 96-well glass bottom plates at a density of 8000-10000 cells per well. Cells were cultured under standard growth conditions for 48 hours prior to FCS measurements. Two distinct controls were maintained and prepared in an identical manner alongside, to establish baseline parameters and account for potential artifacts caused by the fluorophore. Parent control, which is the unmodified, untransfected HeLa Flp-In^TM^ T-Rex cells, seeded to account for the baseline cellular behavior and autofluorescence. EGFP-only control, which is a HeLa cell line with an empty pcDNA5/FRT/TO-EGFP vector, utilized to express and measure diffusion of isolated EGFP-only.

The EGFP-only, EGFP-tagged eIF4B (WT or mutant) and the control parent line cells were maintained in an environmental chamber at 37°C, 5% CO_2_, and controlled humidity. Samples were excited using a 484 nm pulsed diode laser (LDH-D-TA-484, PicoQuant) operating at 20 MHz, with the beam directed through a ZT488/561rpc excitation dichroic filter (Chroma Technology). The emission fluorescence was passed through a 50 µm pinhole conjugate to the sample plane and subsequently split into vertically and horizontally polarized components using a polarizing beam splitter. These polarized signals were filtered through ET535/70 bandpass filters (Chroma Technology) and projected onto two single-photon avalanche diodes (SPADs), which were then auto-correlated eliminating any effect of detector after-pulsing. Data were collected for 2.5 minutes.

For intracellular diffusion measurements, cells were manually selected under brightfield illumination by adjusting the microscope x-y stage. The excitation laser power was carefully optimized to yield a sufficient photon count rate for accurate autocorrelation analysis while preventing photobleaching and phototoxicity. Because EGFP-eIF4B (WT or mutant) localizes exclusively to the cytoplasm, the FCS measurements were targeted to the cytoplasmic compartment by positioning the laser 1 µm above the coverslip interface into the cell. All cell lines, including those for EGFP-only and EGFP-tagged eIF4B (WT or mutant) were maintained in an uninduced state without the addition of tetracycline. The basal (leaky) expression of these constructs provided a strong enough fluorescence signal to generate high-quality autocorrelation curves while maintaining the number of molecules below ∼10 at any given point. Untransfected, parent HeLa cells yielded negligible background auto-fluorescence. Auto-correlation curves were fitted with a diffusion model containing two diffusing species including the EGFP triplet state using Equation 5:

### Mass photometry

All the mass photometry measurements were performed at room temperature using mass photometer (Refeyn, Oxford, UK). The experiments were done using silicon gasket wells (6 individual wells per cassette) that sticks directly on the glass coverslips (#1.5, 24×50 mm^2^, Thermo Fisher). Prior to assembly, the coverslips were cleaned thoroughly with distilled water, ethanol, isopropanol, and subsequently dried. For each measurement, translation buffer alone was used to stabilize and lock the microscope focus prior to sample addition. To convert interferometric contrast to molecular mass, daily calibration curves were generated using proteins and macromolecular standards: erf1 (55 kDa), alcohol dehydrogenase (150 kDa), horse spleen apoferritin (443 kDa), thyroglobulin (669 kDa) and purified ribosomal subunits and complexes (30S, 50S and 70S at 0.9, 1.55, and 2.5 MDa). All calibration standards were diluted to a final concentration of 10 nM. Data were acquired for a duration of 60 seconds. Raw interferometric images were processed and analyzed using DiscoverMP software (Refeyn). Landing events were extracted, and the resulting contrast histograms were fit with a Gaussian function. To determine apparent molecular weights, peak centers were calibrated against a standard curve acquired on the same day.

### Negative stain electron microscopy

Samples of 500 nM eIF4A-eIF4G-eIF4B-WT and eIF4A-eIF4G-eIF4B-mutant along with 0.5 nM dsRNA and 1 mM ATP were prepared. Samples (∼2-3 uL) were adsorbed onto a charged; carbon coated 300 mesh copper grids (Ted Pella Inc.). Grids were then washed with MilliQ water twice, blotting after each application. Washed grids were then stained with 2% uranyl acetate for 1 min and the excess stain was blotted. Imaging was performed on Talos L 120C transmission electron microscopy at 120 kV.

### Recurrence Analysis of Single Particles (RASP)

Single molecule PIE FRET data were acquired as described above, with the exception that the dsRNA concentration was lowered to 0.1 nM to increase the time between the successive single particle fluorescence burst events. Protein concentrations, eIF4A. eIF4G and eIF4B, were kept at the standard values, 0.5 μM each. Data were recorded over a 15 minute interval immediately following ATP addition across multiple independent replicates. Individual 15-minute datasets were converted into .hdf5 files using the ‘phconvert’ module of FRETBursts software and subsequently merged into a single master file for unified RASP processing using a home-build python script. To analyze the probability of a molecule recurring in the confocal volume within a short time window, the FRET efficiency ranges were set to 0.75 – 1.2 and −0.2 – 0.25 for high FRET and low FRET respectively.

## References

1. Linder, P. & Jankowsky, E. From unwinding to clamping- the DEAD-box RNA helicase family. Nat. Rev. Mol. Cell Biol. 12, 505–516 (2011).

2. Özeş, A. R., Feoktistova, K., Avanzino, B. C. & Fraser, C. S. Duplex unwinding and ATPase activities of the DEAD-box helicase eIF4A are coupled by eIF4G and eIF4B. J. Mol. Biol. 412, 674–687 (2011).

3. Rogers, G. W. Jr., Richter, N. J., Lima, W. F. & Merrick, W. C. Modulation of the helicase activity of eIF4A by eIF4B, eIF4H, and eIF4F. J. Biol. Chem. 276, 30914–30922 (2001).

4. Marintchev, A. et al. Topology and regulation of the human eIF4A/4G/4H helicase complex in translation initiation. Cell 136, 447–460 (2009).

5. Oberer, M., Marintchev, A. & Wagner, G. Structural basis for the enhancement of eIF4A helicase activity by eIF4G. Genes Dev. 19, 2212–2223 (2005).

6. Harms, U., Andreou, A. Z., Gubaev, A. & Klostermeier, D. eIF4B, eIF4G and RNA regulate eIF4A activity in translation initiation by modulating the eIF4A conformational cycle. Nucleic Acids Res. 42, 7911–7922 (2014).

7. Izidoro, M. S., Sokabe, M., Villa, N., Merrick, W. C. & Fraser, C. S. Human eukaryotic initiation factor 4E (eIF4E) and the nucleotide-bound state of eIF4A regulate eIF4F binding to RNA. J. Biol. Chem. 298, 102368 (2022).

8. Andreou, A. Z. & Klostermeier, D. The DEAD-box helicase eIF4A: paradigm or the odd one out? RNA Biol. 10, 19–32 (2013).

9. Wolfe, A. L. et al. RNA G-quadruplexes cause eIF4A-dependent oncogene translation in cancer. Nature 513, 65–70 (2014).

10. García-García, C., Frieda, K. L., Feoktistova, K., Fraser, C. S. & Block, S. M. Factor-dependent processivity in human eIF4A DEAD-box helicase. Science 348, 1486–1488 (2015).

11. Sokabe, M. & Fraser, C. S. A helicase-independent activity of eIF4A in promoting mRNA recruitment to the human ribosome. Proc. Natl. Acad. Sci. USA 114, 6304–6309 (2017).

12. Banani, S. F., Lee, H. O., Hyman, A. A. & Rosen, M. K. Biomolecular condensates: organizers of cellular biochemistry. Nat. Rev. Mol. Cell Biol. 18, 285–298 (2017).

13. Guillén-Boixet, J. et al. RNA-induced conformational switching and clustering of G3BP drive stress granule assembly by condensation. Cell 181, 346–361 (2020).

14. Tauber, D. et al. Modulation of RNA condensation by the DEAD-box protein eIF4A. Cell 180, 411–426 (2020).

15. Bohnsack, K. E., Yi, S., Venus, S. & Bohnsack, M. T. Cellular functions of eukaryotic RNA helicases and their links to human diseases. Nat. Rev. Mol. Cell Biol. 24, 749–769 (2023).

16. Choi, J. M., Holehouse, A. S. & Pappu, R. V. Physical principles underlying the complex biology of intracellular phase transitions. Annu. Rev. Biophys. 49, 419–445 (2020).

17. Mittag, T. & Pappu, R. V. A conceptual framework for understanding phase separation and addressing open questions and challenges. Mol. Cell 82, 2201–2214 (2022).

18. King, O. D., Gitler, A. D. & Shorter, J. The tip of the iceberg: RNA-binding proteins with prion-like domains in neurodegenerative disease. Brain Res. 1462, 61–80 (2012).

19. Molliex, A. et al. Phase separation by low complexity domains promotes stress granule assembly and drives pathological fibrillization. Cell 163, 123–133 (2015).

20. Patel, A. et al. A liquid-to-solid phase transition of the ALS protein FUS is promoted by disease mutation and age. Cell 162, 1066–1077 (2015).

21. Wang, J. et al. A molecular grammar governing the driving forces for phase separation of untransformed and complex RNA-binding proteins. Cell 174, 688–699 (2018).

22. Yanas, A., Shweta, H., Owens, M. C., Liu, K. F. & Goldman, Y. E. RNA helicases DDX3X and DDX3Y form nanometer-scale RNA-protein clusters that support catalytic activity. Curr. Biol. 34, 5714–5727 (2024).

23. Nott, T. J. et al. Phase transition of a disordered protein region promotes development of membraneless organelles. Mol. Cell 57, 936–947 (2015).

24. Elbaum-Garfinkle, S. et al. The disordered P granule protein LAF-1 drives phase separation into liquid droplets with tunable viscosity. Proc. Natl. Acad. Sci. USA 112, 7189–7194 (2015).

25. Hondele, A. et al. DEAD-box ATPases are global regulators of phase-separated organelles. Nature 573, 144–148 (2019).

26. Tauber, D., Tauber, G., Khong, A., Van Treeck, B., Pelletier, J., & Parker, R. (2020). Modulation of RNA Condensation by the DEAD-Box Protein eIF4A. Cell, 180(3), 411–426.

27. Swain, B. C. et al. Disordered regions of human eIF4B orchestrate a dynamic self-association landscape. Nat. Commun. 15, 8766 (2024).

28. Feoktistova, K., Tuvshintogs, E., Do, A. & Fraser, C. S. Human eIF4E promotes mRNA restructuring by stimulating eIF4A helicase activity. Proc. Natl. Acad. Sci. USA 110, 13339–13344 (2013).

29. Young, G. et al. Quantitative mass imaging of single biological macromolecules. Science 360, 423–427 (2018).

30. Swaminathan, R., Hoang, C. P. & Verkman, A. S. Photobleaching recovery and anisotropy decay of green fluorescent protein GFP-S65T in solution and cells: cytoplasmic viscosity probed by green fluorescent protein translational and rotational diffusion. Biophys. J. 72, 1900–1907 (1997).

31. Potma, E. O., de Boeij, W. P., Bosgraaf, L., Roelofs, J. & van Haastert, P. J. Reduced protein diffusion rate by cytoskeleton in vegetative and polarized *Dictyostelium* cells. Biophys. J. 81, 2010–2019 (2001).

32. Rogers, G. W. Jr., Lima, W. F. & Merrick, W. C. Biochemical and kinetic characterization of the RNA helicase activity of eukaryotic initiation factor 4A. J. Biol. Chem. 274, 12236–12244 (1999).

33. Andreou, A. Z. & Klostermeier, D. eIF4B and eIF4G jointly stimulate eIF4A ATPase and unwinding activities by modulation of the eIF4A conformational cycle. J. Mol. Biol. 425, 51–61 (2014).

34. Marsden, S., Nardelli, M., Linder, P. & McCarthy, J. E. Unwinding single RNA molecules using helicases involved in eukaryotic translation initiation. J. Mol. Biol. 361, 327–335 (2006).

35. Schmidt, C. et al. eIF4A1-dependent mRNAs employ purine-rich 5′UTR sequences to activate localised eIF4A1-unwinding through eIF4A1-multimerisation to facilitate translation. Nucleic Acids Res. 51, 1859–1879 (2023).

36. Sanders, D. W. et al. Competing protein-RNA interaction networks control multiphase intracellular organization. Cell 181, 306–324 (2020).

37. Conicella, A. E., Zerze, G. H., Mittal, J. & Fawzi, N. L. ALS mutations disrupt phase separation mediated by $\alpha$-helical structure in the TDP-43 low-complexity domain. Structure 24, 1537–1549 (2016).

38. Tsang, B. et al. Phosphoregulated FMRP phase separation models activity-dependent translation through bidirectional control of mRNA granule formation. Proc. Natl. Acad. Sci. USA 116, 4218–4227 (2019).

39. Mitrea, D. M. et al. Nucleophosmin integrates within the nucleolus via multi-modal interactions with proteins displaying R-rich linear motifs and rRNA. eLife 5, e13571 (2016).

40. Kar, M. et al. Solutes unmask differences in clustering versus phase separation of FET proteins. Nat. Commun. 15, 4408 (2024).

