## Supplementary Figures S1 to S7 for "Nanometer-scale RNA protein clusters (RPCs) Foster Helicase Activity of DEAD-box eIF4A"

This file includes:

- Supplementary Figures S1 to S7

**Supplementary Figures and legends:**

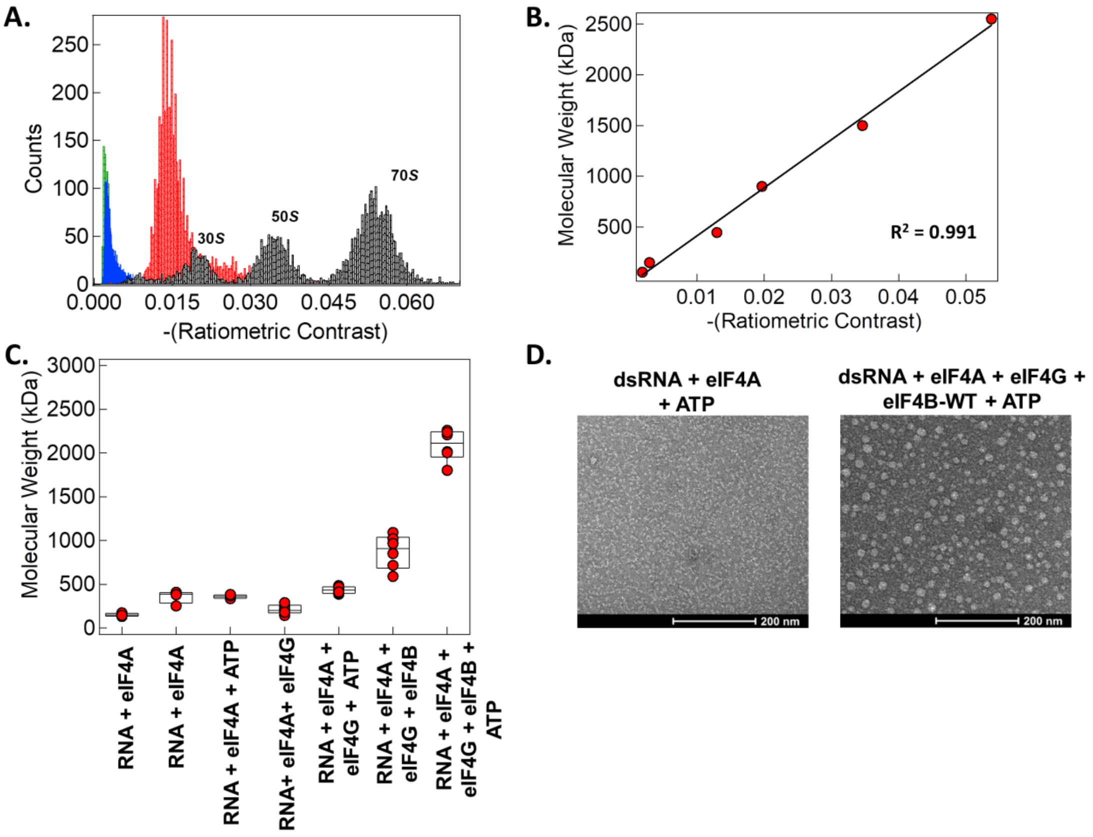


**Figure S1.** (A) Histograms of ratiometric contrasts of eRF1 (green), Alcohol dehydrogenase (blue), Apoferritin (red), and ribosome subunits (30S, 50S and 70S; grey) used as standard to estimate the RPCs size in mass photometry. (B). Molecular mass plotted against ratiometric contrast of standard proteins used for mass calibration. (C) Apparent molecular weights (kDa) of 0.5 μM eIF4A, and eIF4A-eIF4B(WT)-eIF4G with 0.5 nM dsRNA before and after Mg-ATP addition obtained using mass photometry. (D) Representative electron microscopy image of negatively stained eIF4A and eIF4A-eIF4B-eIF4G (0.5 μM each) with 0.5 nM dsRNA plus Mg-ATP. Scale bar, 200 nm.

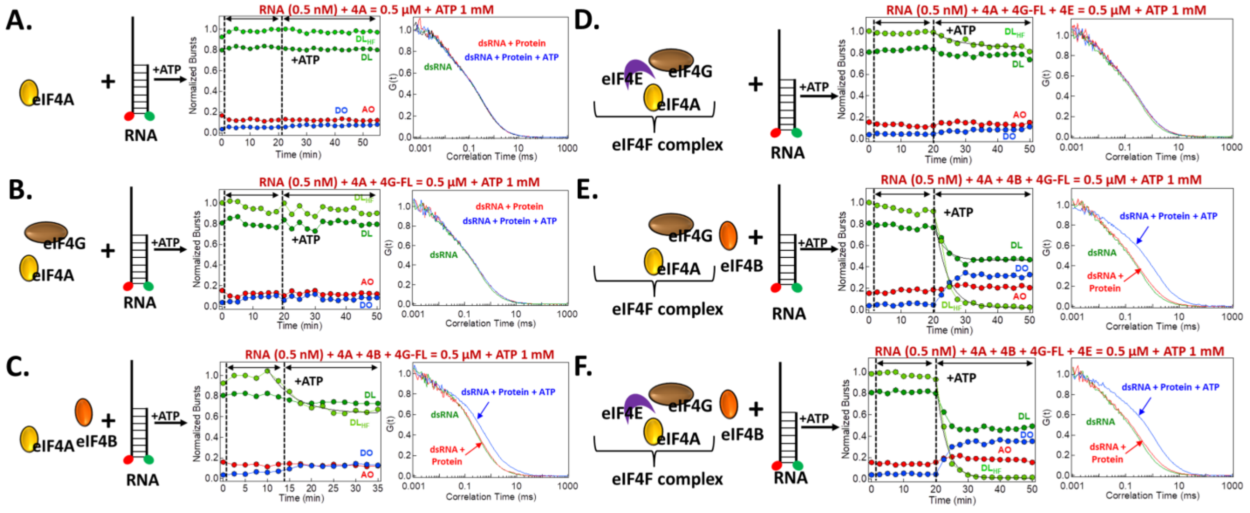


**Figure S2.** Schematic representation of dsRNA and subunits, fraction of DO, DL and AO population along with the normalized auto-correlation curves plotted for (A) dsRNA-eIF4A, (B) dsRNA-eIF4A-eIF4G, (C) dsRNA-eIF4A-eIF4B, (D) dsRNA-eIF4A-eIF4G-eIF4E, (E) dsRNA-eIF4A-eIF4G-eIF4B, and (F) dsRNA-eIF4A-eIF4G-eIF4B-eIF4E for dsRNA only, followed by addition of 0.5 μM protein and subsequent addition of 1mM Mg-ATP. First and second vertical line in the normalized bursts plot represents the time when protein and Mg-ATP were added, respectively.

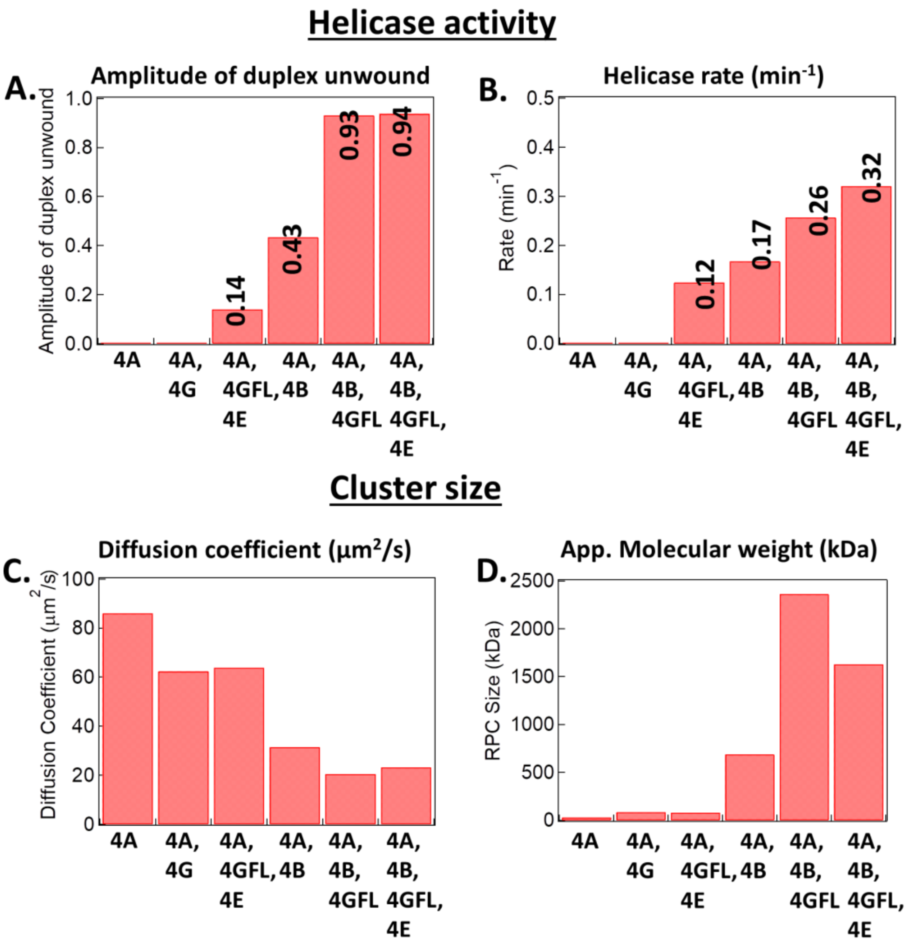


**Figure S3.** Quantification of each subunit on helicase activity in terms of (A) amplitude of duplex unwound (B) helicase rate (min-1) obtained by fitting an exponential to $DL_{HF}$ population, and RPC size in terms of (C) diffusion coefficient ($\mu m^2/s$) and (D) apparent molecular weight (kDa) extracted from FCS data by fitting the auto-correlation curves with diffusion model including triplet. Maximum RNA unwinding activity is obtained when eIF4B is used with eIF4F complex that assembles into large RNA-protein cluster, indicating a central role of eIF4B in both the processes.

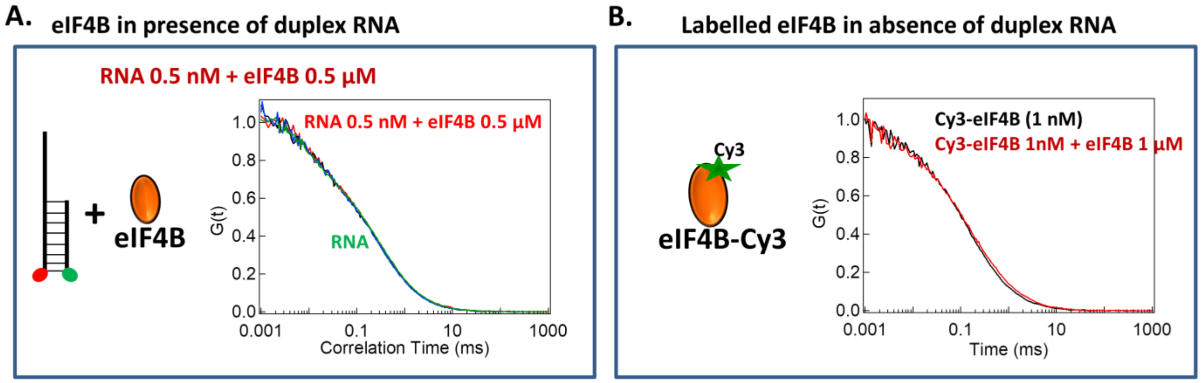


**Figure S4.** (A) Normalized auto-correlation curve of 0.5 nM dsRNA alone (green) and with 0.5 μM eIF4B (red). (B) Normalized auto-correlation curves of 0.5 nM Cy3-eIF4B (black) and with 1 μM unlabeled eIF4B (red). No change in dsRNA or Cy3-eIF4B diffusion is observed with micro-molar concentrations of unlabeled eIF4B.

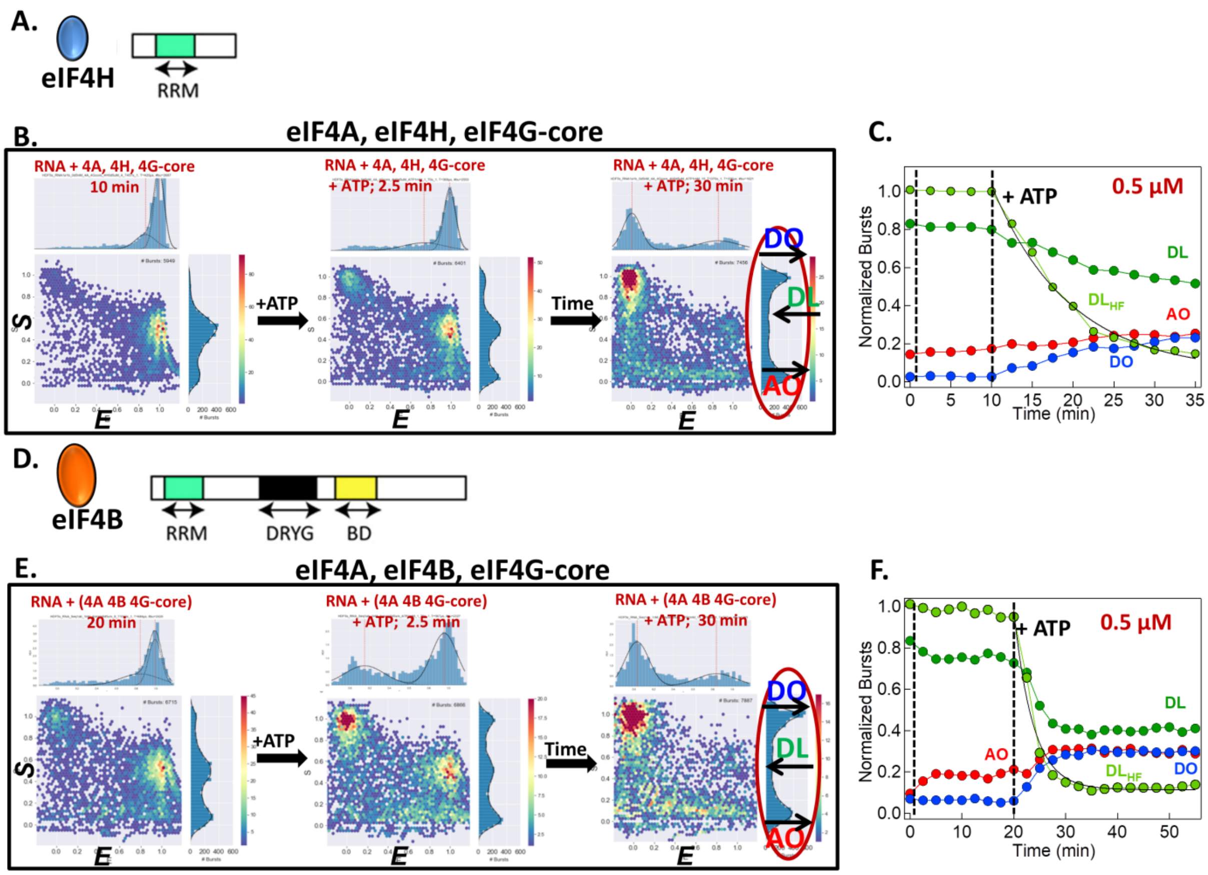


**Figure S5.** (A) Schematic representation of eIF4H, showing the structured RRM region. (B) 2D histogram of *S* vs. *E* showing donor-only (DO, Cy3, *S* = ~1.0), doubly-labelled (DL, *S* = 0.2 – 0.8), and acceptor-only (AO, Alexa647, *S* = ~0.0) population along with the FRET efficiencies (X-axis, *E,* top histogram) and stoichiometry (Y-axis, *S*) histogram (red ellipse). The left scatter plot shows high FRET efficiency, high DL particles, low DO and AO particles for dsRNA with 0.5 μM eIF4A-eIF4H-eIF4G at 10 min before adding Mg-ATP. The middle and right plot shows the three populations after 1 mM Mg-ATP addition at 2.5 min and 30 min. The FRET efficiencies are calculated from DL particles defined as exhibiting *S* in the range 0.25-0.8. (C) The fraction of doubly-labelled (DL; green), doubly-labelled HF($DL_{HF}$; light green), donor-only (DO; blue), and acceptor-only (AO; red) particles are plotted as a function of time for dsRNA alone, following the sequential addition of eIF4A-eIF4H-eIF4G (0.5 μM) and subsequently Mg-ATP (1 mM). First and second vertical line represents the time when protein and Mg-ATP were added, respectively. Helicase rate of 0.105 $min^{-1}$ was extracted by fitting an exponential fit to the $DL_{HF}$. (D) Schematic representation of eIF4B. (E) 2D histogram of *S* vs *E* showing DO, DL and AO population along with FRET efficiencies (X-axis) and stoichiometry (Y-axis) histogram. The left scatter plot shows high FRET efficiency, high DL particles, low DO and AO particles for dsRNA with 0.5 μM eIF4AeIF4B-eIF4G at 10 min before adding Mg-ATP. The middle and right plot shows the three populations after 1 mM Mg-ATP addition at 2.5 min and 30 min. The FRET efficiencies are calculated from DL particles defined as exhibiting *S* in the range 0.25-0.8. (F) The fraction of DL, DO, AO and $DL_{HF}$ particles are plotted as a function of time for dsRNA alone, following the sequential addition of eIF4A-eIF4B-eIF4G (0.5 µM) and subsequently Mg-ATP (1 mM). First and second vertical line represents the time when protein and Mg-ATP were added, respectively.

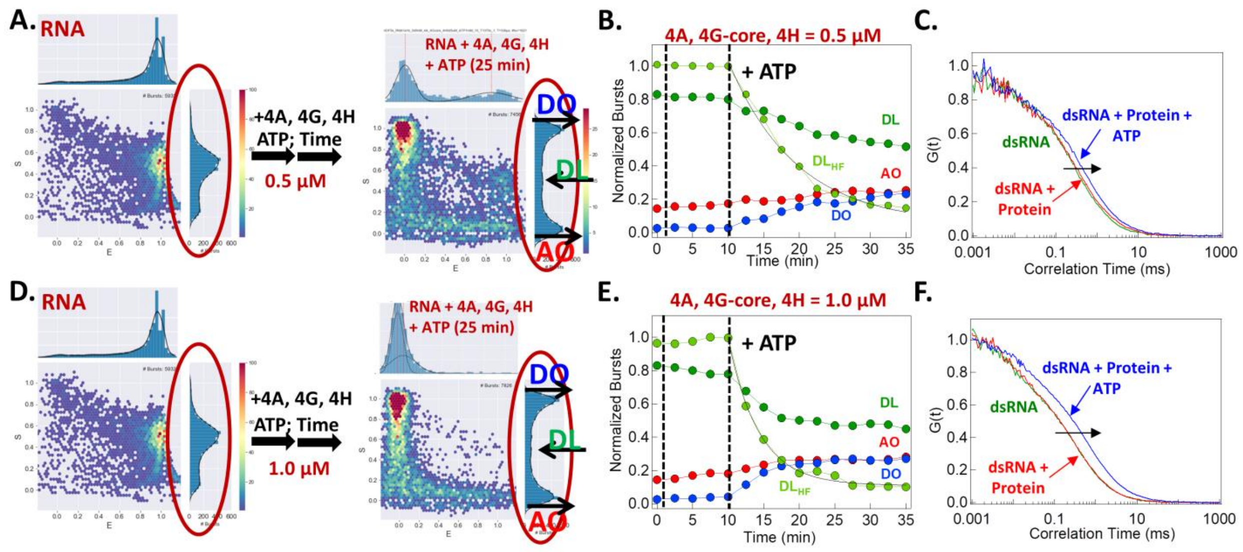


**Figure S6.** (A) 2D histogram of *S* vs. *E* for dsRNA only and with 0.5 μM eIF4A-eIF4H-eIF4G after adding Mg-ATP at 25 min. (B) The fraction of doubly-labelled (DL; green), doubly-labelled HF($DL_{HF}$; light green), donor-only (DO; blue), and acceptor-only (AO; red) particles are plotted as a function of time for dsRNA alone, following the sequential addition of 0.5 μM eIF4A-eIF4H-eIF4G and subsequently Mg-ATP (1 mM). Helicase rate of 0.105 $min^{-1}$ was extracted by fitting an exponential fit to the $DL_{HF}$. (C) Normalized auto-correlation curve for dsRNA (green), after 0.5 μM eIF4A-eIF4H-eIF4G (red), followed by the addition of Mg-ATP (blue). A diffusion coefficient of 42.66 ± 2.72 $μm^2/s$ (mean ± s.e.m., n=3) was obtained by fitting the acceptor auto-correlation after Mg-ATP addition. (D) 2D histogram of *S* vs. *E* for dsRNA only and with 1 μM eIF4A-eIF4H-eIF4G at 25 min after adding Mg-ATP. (E) The fraction of DL, $DL_{HF}$, DO and AO particles are plotted as a function of time for dsRNA alone, following the sequential addition of eIF4A-eIF4H-eIF4G (1 μM) and subsequently Mg-ATP (1 mM). Helicase rate of 0.223 $min^{-1}$ was extracted by fitting an exponential fit to the $DL_{HF}$. (F) Normalized auto-correlation curve for dsRNA (green), after 1 μM eIF4A-eIF4H-eIF4G (red), followed by the addition of Mg-ATP (blue). A diffusion coefficient of 33.3 ± 1.76 $μm^2/s$ (mean ± s.e.m., n=3) was obtained by fitting the acceptor auto-correlation after Mg-ATP addition. Both, the RPC size and helicase rate showed concentration dependent increase with increasing eIF4H concentration.

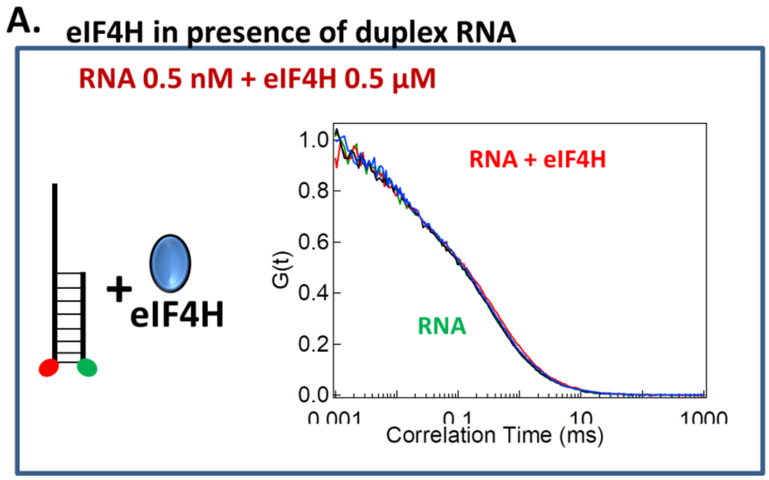


**Figure S7.** (A) Normalized auto-correlation curve of 0.5 nM dsRNA alone (green) and with 0.5 µM eIF4H (red). No change in dsRNA diffusion is observed with micro-molar concentrations of unlabeled eIF4H.
